# Long-term population studies uncover the genome structure and genetic basis of xenobiotic and host plant adaptation in the herbivore *Tetranychus urticae*

**DOI:** 10.1101/474064

**Authors:** Nicky Wybouw, Olivia Kosterlitz, Andre H. Kurlovs, Sabina Bajda, Robert Greenhalgh, Simon Snoeck, Huyen Bui, Astrid Bryon, Wannes Dermauw, Thomas Van Leeuwen, Richard M. Clark

## Abstract

Pesticide resistance arises rapidly in arthropod herbivores, as can host plant adaptation, and both are significant problems in agriculture. These traits have been challenging to study as both are often polygenic and many arthropods are genetically intractable. Here, we examined the genetic architecture of pesticide resistance and host plant adaptation in the two-spotted spider mite, *Tetranychus urticae,* a global agricultural pest. We show that the short generation time and high fecundity of *T. urticae* can be readily exploited in experimental evolution designs for high-resolution mapping of quantitative traits. As revealed by selection with spirodiclofen, an acetyl-CoA decarboxylase inhibitor, in populations from a cross between a spirodiclofen resistant and a susceptible strain, and which also differed in performance on tomato, we found that a limited number of loci could explain quantitative resistance to this compound. These were resolved to narrow genomic intervals, suggesting specific candidate genes, including *acetyl-CoA decarboxylase* itself, clustered and copy variable cytochrome P450 genes, and *NADPH cytochrome P450 reductase*, which encodes a redox partner for cytochrome P450s. For performance on tomato, candidate genomic regions for response to selection were distinct from those responding to the synthetic compound and were consistent with a more polygenic architecture. In accomplishing this work, we exploited the continuous nature of allele frequency changes across experimental populations to resolve the existing fragmented *T.urticae* draft genome to pseudochromosomes. This improved assembly was indispensable for our analyses, as it will be for future research with this model herbivore that is exceptionally amenable to genetic studies.

## INTRODUCTION

Although pesticides with diverse modes of action have been developed to combat populations of insect and mite herbivores, the evolution of resistance is common. As early as 1937, Theodosius Dobzhansky noted that the emergence of resistance to chemical pesticides in insect populations was “probably the best proof of the effectiveness of natural selection yet obtained” (Dobzhansky 1937; Ceccatti 2009). In the intervening years, numerous studies have implicated genetic variants in the molecular targets of pesticides as underlying “target-site” resistance. A second major route to resistance involves genetic changes that affect penetration, metabolism, sequestration and excretion of pesticides (toxicokinetic resistance) (Feyereisen *et al.* 2015). Of these, metabolic mechanisms have been especially well studied, and genetic changes affecting the coding sequences and transcription of genes in detoxification families, like cytochrome P450 monooxygenases (CYPs) and carboxyl/cholinesterases (CCEs), have been implicated in the metabolism of xenobiotics in diverse organisms (Li *et al.* 2007; Feyereisen *et al.* 2015; Van Leeuwen and Dermauw 2016).

Despite the ubiquity of pesticide resistance across arthropod species (Sparks and Nauen 2015), as well as progress in understanding the molecular mechanisms of toxicokinetic processes, questions about the genetic architecture and evolutionary origins of pesticide resistance remain (Hawkins *et al.* 2018). Numerous studies have shown that the genetic architecture of resistance in herbivores can be variable (ffrench-Constant *et al.* 2004; Van Leeuwen *et al.* 2010; Feyereisen *et al.* 2015). In some cases, a monogenic change, typically in a target-site, leads to high resistance levels observed in field populations (Roush and McKenzie 1987; Van Leeuwen *et al.* 2008, 2012; Douris *et al.* 2016; Riga *et al.* 2017). Nevertheless, high-level resistance to pesticides in herbivore populations is often polygenic, and in most cases the number of causal loci, their relative effect sizes, the nature of the underlying loci and alleles, and their origins, are unknown (Li *et al.* 2007; Hawkins *et al.* 2018). More generally, detailed understandings of the genetic architecture of resistance in arthropods come disproportionally from insects like *Drosophila melanogaster* or mosquito species, for which discovery of resistance loci has been facilitated by dense genetic and genomic resources (ffrench-Constant *et al.* 2004; Hemingway *et al.* 2004; Ranson *et al.* 2004). In contrast, for most herbivores, even major global pests, these resources are minimal or absent. In addition, the life histories or breeding systems of many herbivores hamper genetic approaches. Although large-effect quantitative trait loci (QTL) for resistance have been mapped in some arthropod herbivores, they frequently encompass large chromosomal regions (Gahan 2001; Saavedra-Rodriguez *et al.* 2008; Coates and Siegfried 2015; Coates *et al.* 2016).

Therefore, inferences about mechanisms of pesticide resistance in herbivore populations have often come from other approaches. For instance, expression studies have frequently been employed to identify genes induced or constitutively overexpressed in pesticide resistant strains (Oppenheim *et al.* 2015). Where resulting candidate genes are amenable to functional assays, as for CYPs and CCEs, enzymatic modification of pesticides *in vitro* has often been taken to suggest causality. Nonetheless, whether such candidates contribute to resistance *in vivo*, and their relative contribution in the case of polygenic resistance, is generally not known. Further, expression studies typically identify hundreds of candidate genes, many of which have unknown functions (Grbić *et al.* 2011; Dermauw *et al.* 2013; Bansal *et al.* 2014), or alternatively belong to gene families for which heterologous assays are either challenging or not established. The skewed focus on genes in a small number of experimentally tractable detoxification families has therefore potentially led to a biased view of the spectra of loci that contribute to pesticide resistance.

Mirroring long-standing interest in the evolution of pesticide resistance in herbivores, the genetic basis of the evolution of host plant use has attracted intense interest, as has the question of whether the latter may facilitate the former (Dermauw *et al.* 2018; Hardy *et al.*2018). Typically, an herbivore encounters a set of defensive compounds in its diet. This is especially true for generalist herbivores, which encounter different blends of toxins across their host plant ranges (Strong *et al.* 1984; Schoonhoven *et al.* 2005). As observed for pesticide resistance, traditional genetic mapping studies using microsatellites and other genetic markers have revealed that the genetic architecture of host plant use is variable, and ranges from oligogenic to polygenic (Jones 1998; Midamegbe *et al.* 2011; Henniges-Janssen *et al.* 2011; Jaquiéry *et al.* 2012; Oppenheim *et al.* 2012; Alexandre *et al.* 2013; Nouhaud *et al.* 2014). However, with few exceptions (Bass *et al.* 2013; Wybouw *et al.* 2014), candidate genes for host plant adaptation are unknown, as is whether loci for host plant use are also targets of selection for resistance to synthetic pesticides.

In the current study, we examined the genetic basis of pesticide resistance and plant host use in the two-spotted spider mite, *Tetranychus urticae* (Arthropoda: Chelicerata: Acari: Acariformes). This herbivore is a globally distributed agricultural pest, and has among the highest documented occurrences of pesticide resistance (Van Leeuwen *et al.* 2010). *T. urticae* is also an extreme generalist (Migeon *et al.* 2010), and numerous experimental studies have shown that populations can adapt quickly to new host plants (Gould 1979; Fry 1989; Agrawal 2000; Magalhães *et al.* 2007, 2009; Wybouw *et al.* 2015). Among herbivores, this species is exceptionally tractable for genetic and genomic studies. *T. urticae* has a fast generation time – less than ten days at high temperatures – and can be maintained in laboratory populations of thousands on single plants (Grbić *et al.* 2007; Van Leeuwen and Dermauw 2016). Prior cytological work has suggested that the species has three holocentric chromosomes (Helle and Bolland 1967; Grbić *et al.* 2007), and the genome is compact, with a draft Sanger assembly having a cumulative length of only 90.8 Mb (Grbić *et al.* 2011). Further, annotation and gene expression studies revealed expansions of detoxification gene families, as well as gene families that change in expression upon pesticide exposure or host plant shift, but that had not been previously associated with adaptation to either (Grbić *et al.* 2011; Dermauw *et al.* 2013; Wybouw *et al.* 2015; Snoeck *et al.* 2018).

To start, we generated an inbred strain, SR-VP, from a field collected *T. urticae* population resistant to spirodiclofen, a recent but widely used compound that disrupts lipid synthesis by inhibiting acetyl-CoA decarboxylase (ACCase) (Bretschneider *et al.* 2007; Lümmen *et al.* 2014). Prior genetic studies with the parental population were consistent with polygenic resistance, and a combination of approaches, including gene expression studies and *in vitro* tests for spirodiclofen metabolism by several candidate CYPs, suggested a role for metabolic resistance (Van Pottelberge *et al.* 2009; Demaeght *et al.* 2013). We found that the SR-VP strain retained high-level resistance to spirodiclofen; additionally, it performed an order of magnitude better on tomato (*Solanum lycopersicum*), a challenging host for many spider mite populations (Agrawal *et al.* 2002; Wybouw *et al.* 2015), than did a spirodiclofen-susceptible inbred strain, Lon-Inb.

With large, replicated populations from a SR-VP × Lon-Inb cross that were maintained with or without selection for ~50 generations, we identified genomic regions responding to both spirodiclofen treatment and growth on tomato. This allowed us to address questions about the number of loci that underpin pesticide resistance, the genetic architecture of host plant adaptation, and the relationship between them. In some cases, we localized QTL to narrow genomic intervals that highlighted specific candidate genes and alleles. To accomplish this work, we exploited the continuous nature of allele frequency changes in experimental populations to consolidate the fragmented *T. urticae* draft genome into a chromosome-level assembly. This resource enabled our characterization of genome-wide responses to selection, as it will for future studies with this experimentally tractable herbivore.

## MATERIALS AND METHODS

### Biological materials

Genetic crosses and phenotypic selections were performed with two inbred *T. urticae* strains, Lon-Inb and SR-VP. Strain London, which was used to construct the reference Sanger draft assembly for *T. urticae* (hereafter the Sanger assembly) (Grbić *et al.* 2011), was not initially inbred (Van Leeuwen *et al.* 2012). Subsequently, it was inbred by seven rounds of mother-son crosses to produce Lon-Inb (Díaz-Riquelme *et al.* 2016). Starting with a previously reported spirodiclofen-resistant population, SR-VP (Van Pottelberge *et al.* 2009; Demaeght *et al.* 2013), we performed six generations of mother-son crosses to create the SR-VP inbred strain (hereafter simply SR-VP; inbreeding was performed as described by Bryon *et al.* 2017a). To facilitate genomic analyses, six additional *T. urticae* strains were collected as part of this study (Heber, Parrott, RS, ShCo, TT, and WG-Del; generations of inbreeding is given in Table S1).

### Experimental design for host plant and pesticide selections

The Lon-Inb and SR-VP strains were kept on potted bean plants (*Phaseolus vulgaris*) under laboratory conditions (25°C, 60% relative humidity and 16:8 hour light:dark photoperiod). An F1 hybrid population was generated by crossing 1500 one-day-old virgin adult Lon-Inb females with 600 one-to two-day-old adult SR-VP males. We allowed the hybrid population to expand for approximately five generations, and from this established 15 populations on potted bean plants (unselected controls) and 15 populations on tomato plants (tomato selections; *S. lycopersicum* cv Moneymaker). These populations were founded with 500 adult females. Subsequently, from each of the 15 control populations on bean, a paired spirodiclofen selection population was established by transferring approximately 1000 mites to bean plants that were sprayed with 100 mg/L spirodiclofen (commercial formulation, Envidor^®^ 240 g/L SC, Bayer Crop Science, Leverkusen, Germany). Mites from these 45 populations were propagated for over 50 generations. New three-week-old tomato and bean plants (spirodiclofen-sprayed and non-sprayed) were offered to the respective selection and control populations on a weekly basis. During the course of the experiment, the selection pressure of spirodiclofen was gradually increased until no acaricide related mortality was observed on beans sprayed until run-off with 5000 mg/L of spirodiclofen.

### Phenotypic analyses of evolved populations

Mites from the ancestral strains and derived populations across the three treatments (tomato selection, spirodiclofen selection and control) were reared on unsprayed bean plants for two generations to remove acclimation or maternal effects. Experimental evolution to spirodiclofen was evaluated by performing larvicidal toxicity bioassays as previously described (Van Pottelberge *et al.* 2009). Briefly, leaf discs were sprayed in an Auto Loading Potter Lab spray tower (Burkard Scientific, Uxbridge, UK) with settings of 1 bar and 850 µl with a 0.002 g/cm^2^coverage. Survival was scored at the deutonymphal stage. The spirodiclofen concentrations lethal to half the population (LC_50_ values) of the parental strains were estimated using probit analysis (POLO; LeOra Software, Berkeley, CA). For the parental SR-VP and Lon-Inb strains, differences in survival percentages at 2500 mg/L and 5000 mg/L spirodiclofen were assessed using a generalized linear model with a binomial distribution (proc genmod in SAS, version 9.4, SAS Institute, Cary, NC) with strain and dosage as fixed effects. Survival percentages between the paired spirodiclofen-selected and control populations were analyzed using a generalized linear mixed model with a binomial distribution (proc glimmix in SAS). Here, selection regime and dosage were incorporated as fixed effects in the linear model, whereas population was regarded as a random effect.

Experimental evolution on tomato was evaluated by transferring 35 two-day-old females to the three leaflets of a fully developed tomato leaf with four replicates per population. Performance was estimated ten days post-infestation by measuring total mite population sizes on the respective tomato plants (Wybouw *et al.* 2015). Performance of the two parental strains on tomato was analyzed using a general linear model (proc glm in SAS). Differences in mite performance on tomato between the tomato-selected and control populations were assessed by a general linear mixed model with the selection regime and populations as fixed and random effects, respectively (proc mixed in SAS).

### Genome sequencing

For *T. urticae* strains, genomic DNA preparation and quality assessment, construction of Illumina libraries, and sequencing at either the Centro Nacional de Análisis Genómico (CNAG, Barcelona, Spain) or the High-Throughput Genomics and Bioinformatic Analysis Shared Resource at the Huntsman Cancer Institute (University of Utah, Salt Lake City, UT) was as previously described (Bryon *et al.* 2017a). For Lon-Inb and SR-VP paired-end reads of 101 bp were generated, and for all other strains except TT and RS, paired-end 125 bp reads were produced; for TT, single-end 50 bp reads were generated, while for RS, paired-end 300 bp reads were produced on an Illumina MiSeq instrument (Table S1).

To analyze the impact of spirodiclofen selection on genome-wide allele frequencies, we prepared DNA from eight spirodiclofen-selected populations that responded strongly to selection, along with their matching unselected control populations (Figure 1A). We also prepared DNA from five tomato-selected and bean control populations with respectively high and low performance on tomato (Figure 1B). For each of the resulting 22 experimental populations (some control populations were shared between selections, Figure 1A,B), genomic DNA was extracted from ~800 to 1000 pooled adult females collected at the end of the selection experiments using a phenol-chloroform method as previously described (Van Leeuwen *et al.* 2012) and washed twice with 70% ethanol. DNA samples were subsequently purified with an EZNA Cycle Pure Kit (Omega Bio-tek, VWR, Amsterdam, the Netherlands) and eluted in 35 µl of TE buffer provided by the purification kit. DNA concentrations and integrity were measured with a Qubit (Thermo Fisher Scientific, Waltham, MA) and an Agilent TapeStation 2200 (Software A.01.04; Agilent Technologies, Santa Clara, CA), respectively. For each population sample, paired-end reads of 125 bp in length were generated at the High-Throughput Genomics and Bioinformatic Analysis Shared Resource.

**Figure 1.**
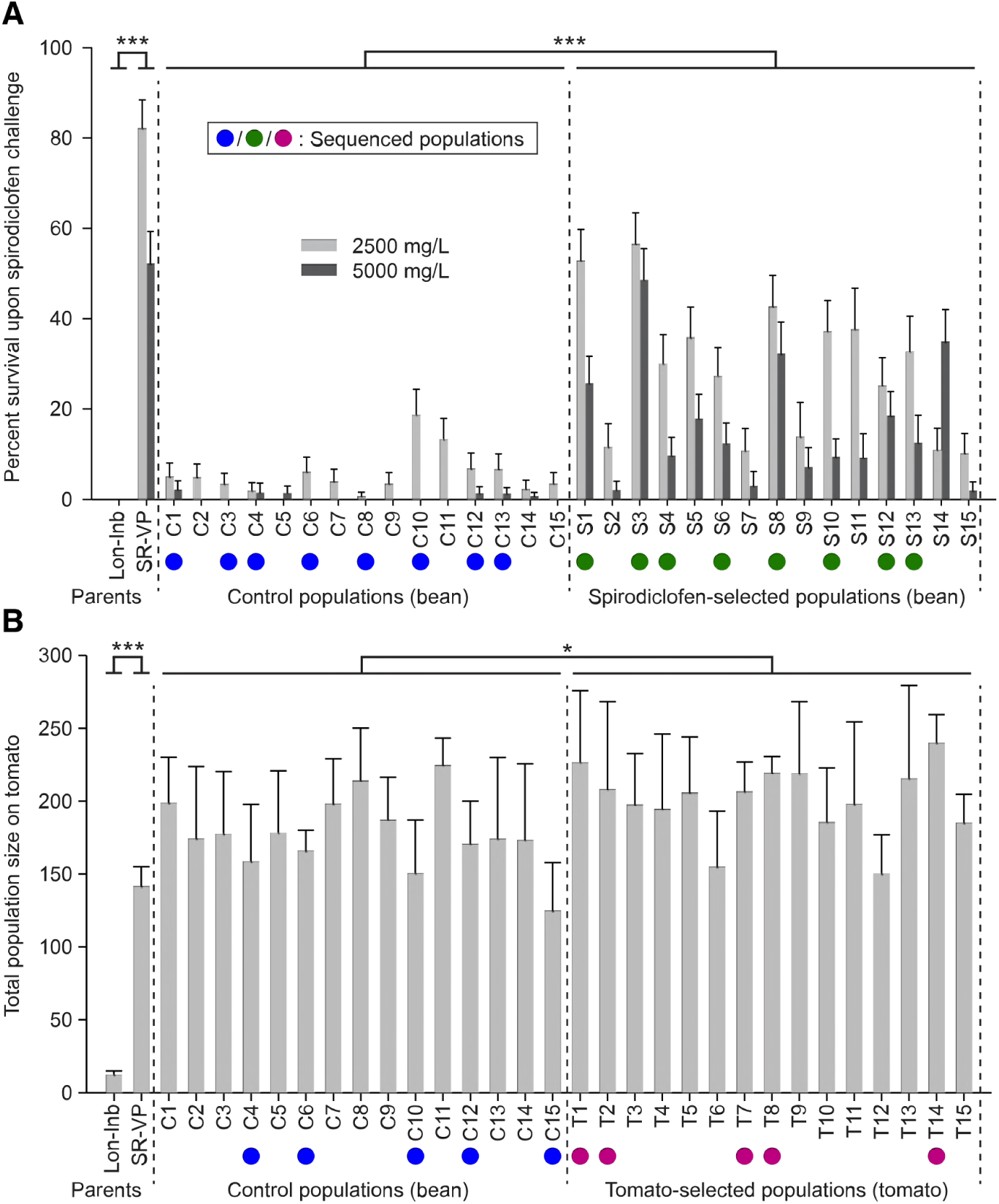
Response to selection in parental strains and experimental populations. (A) Survival at the deutonymphal stage after spraying with 2500 and 5000 mg/L of spirodiclofen (Envidor). Differences in survival were present between the parental strains (Lon-Inb and SR-VP) and spirodiclofen-selected and control populations as indicated. (B) Performance on tomato as assessed by total population size 10 days after initial plant inoculation with 35 founding females. Significant differences between the parental strains and tomato-selected and control populations are as indicated. Populations selected for genomic sequencing are indicated by colored circles: blue, control populations; green, populations grown on bean, selected with spirodiclofen; purple, populations maintained on tomato. In all plots, bars represent two standard errors of the mean. Statistical significance: *p* < 0.05, *; *p* < 0.0001, ***.

### Identification of high-quality variants

Reads from *T. urticae* strains and experimental populations (Table S1) were aligned to the *T. urticae* Sanger assembly (Grbić *et al.* 2011) using the default settings of BWA-MEM 0.7.15-r1140 (Li 2013), and were sorted by coordinate using SAMtools 1.3.1 (Li *et al.* 2009). Duplicate reads were subsequently marked with Picard 2.6.0 (http://broadinstitute.github.io/picard) prior to indel realignment with GATK 3.6-0-g89b7209 (Van der Auwera *et al.* 2013). Across Lon-Inb, SR-VP, and the 22 experimental populations, single nucleotide polymorphisms (SNPs) and indels were called using the GATK 3.6-0-g89b7209 UnifiedGenotyper tool; the output of this analysis was a single Variant Call Format (VCF) file from which allele frequencies at variable sites were extracted for downstream analyses. We also performed a similar analysis with short-read data from two prior bulked segregant analysis (BSA) genetic mapping studies in *T. urticae* (Van Leeuwen *et al.* 2012; Bryon *et al.* 2017a) using the same read mapping and variant prediction workflow. To select high-quality SNPs in the respective data sets (parental strains and derived populations), we parsed the VCF files to identify alternative alleles that were fixed but different in parental strains and had Phred-scaled quality scores >100. Additionally, on a per sample basis, we required that read coverage at a variable site be within 25-150% of the respective sample’s genome-wide mean as assessed at all variable positions. For the Bryon *et al*. (2017a) data, the second parent could not be inferred directly as single males founded crosses (male *T. urticae* are haploid). For inference, we adopted the same methods used in that study (Bryon *et al.* 2017a).

### Principal component analysis

For the control, spirodiclofen-selected, and tomato-selected populations, a principal component analysis (PCA) was performed using a correlation matrix of SNP frequencies as extracted from the respective VCF file (R function prcomp, which is part of the R-package ‘stats’, version 3.3.0) (R Core Team 2016). To be included in the PCA (Figure 2), variable positions had to pass the filters we used to select high-quality SNPs in each of the 22 experimental samples. The PCA plot was created with autoplot, a function of the R-package ‘ggplot2’ version 2.1.0 (Wickham 2016).

**Figure 2.**
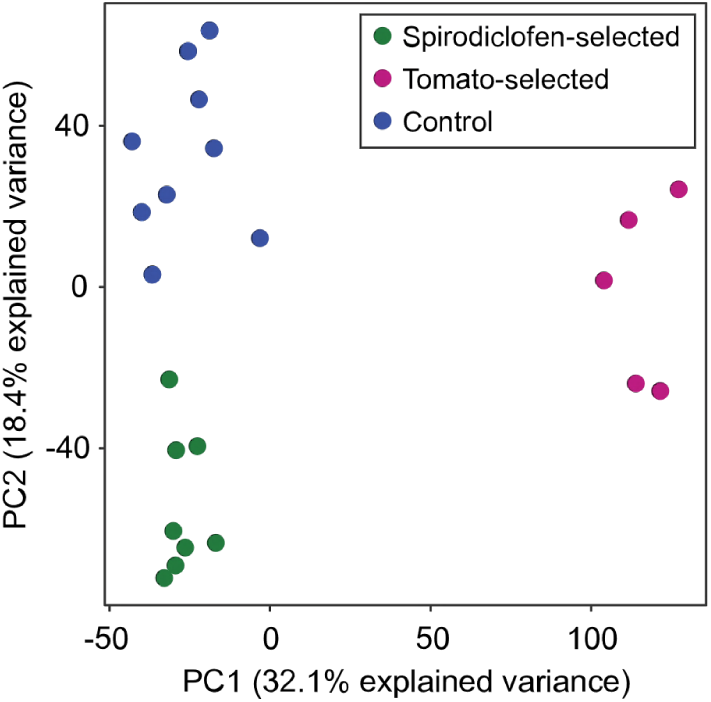
Genomic responses to selection by spirodiclofen and tomato differ. Principal component analysis of control, spirodiclofen-selected, and tomato-selected populations with genome-wide allele frequency data at SNP loci. Circles represent individual populations colored by treatment as indicated (legend, top right; compare to Figure 1).

### *De novo* assemblies of seven *T. urticae* strains

We generated *de novo* assemblies for inbred *T. urticae* strains (Table S1) and aligned them to the Sanger assembly. Illumina reads were imported into CLC Genomics Workbench 9.0.1 (https://www.qiagenbioinformatics.com/) and trimmed using the following settings: quality score limit of 0.05 and maximum of 2 ambiguous nucleotides. *De novo* assemblies for each strain were subsequently constructed with the short-read data using the following options: Automatic word and bubble size, Minimum contig length of 200, Auto detection of paired distances, Perform scaffolding, and Map reads back to contigs (Mismatch cost: 2, Insertion cost: 3, Deletion cost: 3, Length fraction: 0.5, Similarity fraction 0.8, and Update contigs checked). The resulting sequences for each strain were aligned to the Sanger assembly using the default settings of the BLASR 1.3.1 aligner (Chaisson and Tesler 2012). The alignments were subsequently converted to coordinate-sorted and indexed BAM files using SAMtools 1.3.1 (Li *et al.* 2009).

### Identification of misassembled regions in Sanger scaffolds

To locate potential misassemblies in the Sanger reference sequence, we identified abrupt shifts in allele frequencies in experimental populations as a function of genomic position. To do this, we developed a metric, the average window distance (AWD), for which large values between adjacent (non-overlapping) genomic windows are expected to reflect Sanger assembly errors. A schematic illustrating the AWD metric, and the principle behind its use to detect misassembled regions, is given in Figure 3. Briefly, for our study we computed AWD values with 10 kb offsets for all informative, immediately adjacent non-overlapping 150 kb windows across concatenated Sanger scaffolds (ordered by decreasing length). For this analysis, we included scaffolds 1-44 as they harbor ~95% of the cumulative Sanger assembly length, and were included in three prior BSA mapping studies in *T. urticae* (Van Leeuwen *et al.* 2012; Demaeght *et al.* 2014; Bryon *et al.* 2017a). The remaining 596 scaffolds decrease rapidly in length, with a median of only 4 kb, are often repetitive, and contain sequences potentially allelic to those found in the larger scaffolds (Grbić *et al.* 2011; Wybouw *et al.* 2018). For windows to be informative across all samples (experimental populations and the two parents), they had to have ≥20 high quality SNPs; if not, the nearest adjacent windows meeting these criteria were used. To compute AWD values from informative non-overlapping windows, we: (1) determined each window’s SR-VP allele frequency (as assessed from all aligned reads at informative SNP loci in the windows) for each of the 22 study populations individually, (2) calculated the absolute values of the differences in the two frequency values between the non-overlapping windows on a per population basis, and (3) averaged the resulting 22 values (see Figure 3 for example calculations).

**Figure 3.**
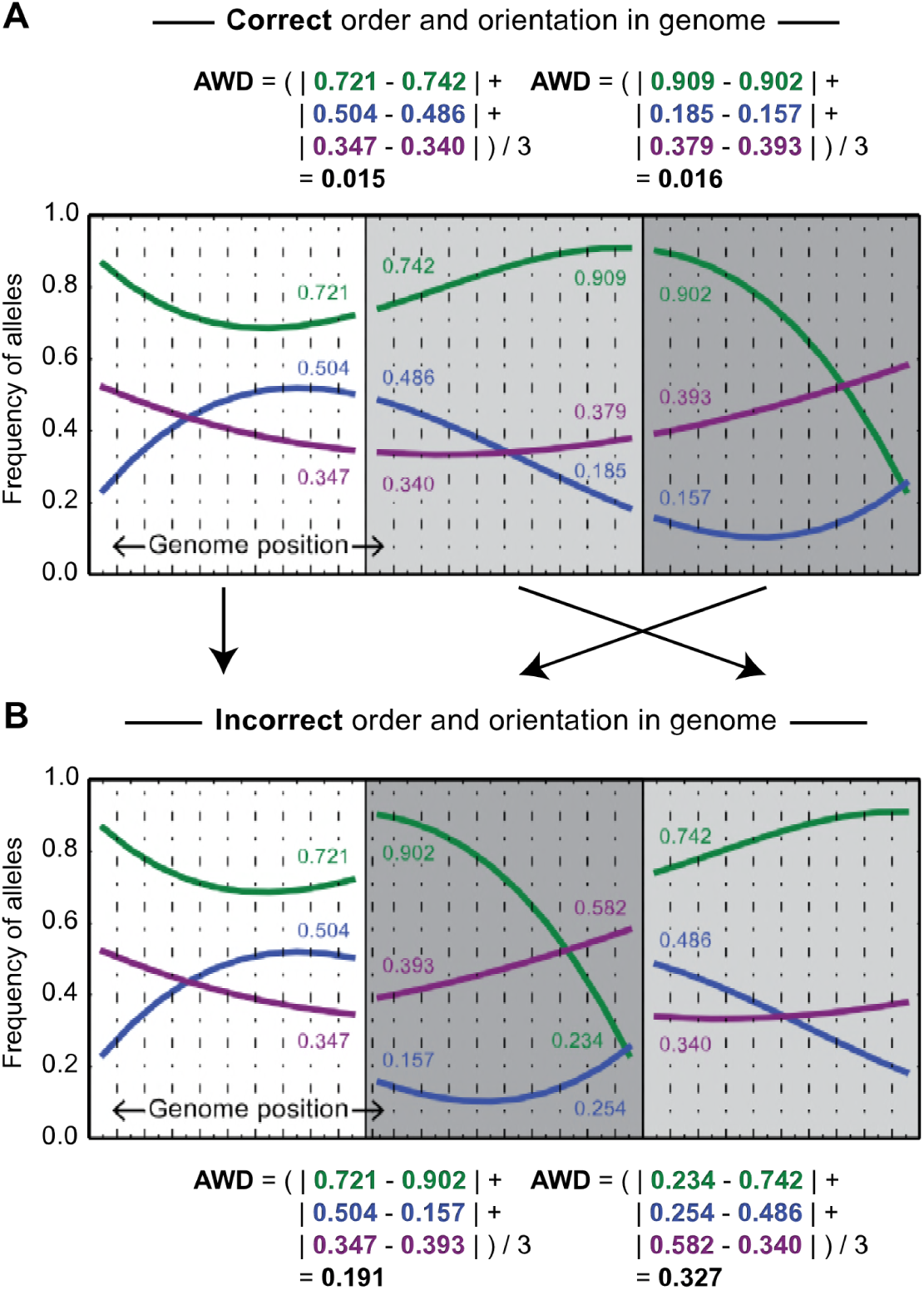
Illustration of the average window distance (AWD) metric used for genome curation and assembly. (A) From experimental populations, allele frequencies, which can be estimated from short-read data (e.g., Figure 4 and Figures S1, S3 and S4), change proportionally to distance along chromosomes due to genetic drift or potentially selection. A schematic of a plausible pattern for three populations is shown (colored lines). At given sites in the genome (solid vertical lines with different intensity of shading on either side), the difference in allele frequencies in adjacent windows on either side of the solid lines will approach zero. At top, AWD calculations are shown for a genomic region in which the assembly is correct. (B) The same schematic as shown in panel A except that the two shaded genomic regions have been shuffled (that is, they are out of their true order). Now, when AWD values are calculated (bottom), the values are elevated markedly above zero. This illustrates how anomalously high AWD values detect misassemblies (Figure 4A); in a related approach, for unordered genomic regions (e.g., scaffolds in draft genomes), minimal AWD values between pairs of scaffolds suggest adjacency and orientation (Figure 4).

To investigate potential misassemblies suggested by high AWD values, we then examined peaks greater than two standard deviations from the genome-wide AWD mean (0.07). Potential sources of elevated “blips” in AWD values, other than true assembly errors, are small transpositions, or simply errant SNP predictions in complex, local regions. Therefore, we removed windows yielding AWD values >0.07 (the masked regions spanned all pairs of windows that yielded above-threshold AWD values), and then recalculated AWD values across the genome. Excluding junctions between scaffolds, which are *de facto* misassemblies in the concatenated genome (Figure 1), regions of elevated AWD values that did not drop below 0.07 were investigated further (Figure S1). To do this, we examined alignments of short-read *de novo* assemblies from seven *T. urticae* strains (Table S1) to the Sanger reference sequence in the regions of elevated AWD values. BLASR 1.3.1, which was used to generate the alignments, allows contigs (or portions thereof) to be aligned to multiple genomic locations. If one set of aligned short-read assembled contigs (or contig portions) ended, and another set started within the same 5 bp at the site of an anomalously high AWD value internal to a Sanger scaffold, we considered the location a misassembly. For these instances, scaffolds were broken into subscaffolds for subsequent analyses (four out of six instances, on scaffolds 1, 2, 4, and 8 as indicated in Table S2). For the remaining cases (Figure S1), short-read *de novo* assemblies either did not suggest misassemblies, or were uninformative (no *de novo* contigs aligned at the sites of the elevated AWD values).

### Construction of superscaffolds with population allele frequency data

To place and order Sanger scaffolds relative to one another, we chained Sanger scaffolds (including subscaffolds) together based on reciprocal lowest AWD values as calculated from terminal windows on the scaffolds. All Sanger scaffolds and subscaffolds of at least 100 kb in length were used in the process (Sanger scaffolds 1-44), except for scaffold 42, which was excluded because of extreme copy variation (Figure S2). AWD calculations were as for misassembly detection, except only terminal windows were used, hereafter called scaffold segment ends (SSEs). SSEs lengths were 300 kb, but were offset 50 kb internally to the ends of Sanger scaffolds; this offset was used to avoid potentially repetitive sequences at the ends of Sanger scaffolds (i.e., we deemed that variant detection at Sanger scaffold ends might be unreliable). If a Sanger scaffold was shorter than the window length plus the offsets (400 kb in total, corresponding to Sanger scaffolds 39 and higher), the entire Sanger scaffold was treated as a single SSE (i.e., one non-oriented window).

With the resulting 94 SSEs, AWDs were then calculated among all pairs. For each SSE, a list was produced with all non-self AWD comparisons containing the SSE (a total of 93 comparisons) and sorted in ascending order. The five smallest AWD comparisons were retained for downstream analyses; in this scheme, a small AWD value supports proximity in the genome (Figure 3). For two SSEs, reciprocal smallest AWD values were taken as evidence of adjacency and relative orientation. When this occurred, the two SSEs were removed from all SSE lists and excluded from subsequent analyses, and the process was repeated iteratively with unmatched SSEs until there were no more rankings to compare (in this process, reciprocal best hits of two SSEs on the same scaffold were ignored). An exception was for Sanger scaffolds 39 and greater; as only a single SSE could be calculated for these small scaffolds, they were only removed once they had been called twice (this allowed these Sanger scaffolds to potentially connect to other Sanger scaffolds on both sides). Sanger scaffold endings with unpaired SSEs, a result of no reciprocal AWD matches in the initial top five rankings in the terminal iteration, were considered as putative chromosome ends and were used to initiate the construction of superscaffolds by placing and ordering Sanger scaffolds according to the catalog of reciprocal best hits. An exception was for Sanger scaffolds less than 400 kb; because these were treated as single windows, their forward or reverse orientations in resulting superscaffolds could not be determined, even though they could be placed between flanking Sanger scaffolds.

### Construction of pseudochromosomes by incorporating *de novo* assembly data

As a complementary method for ordering and orienting scaffolds in the Sanger assembly, and to validate the AWD-based joining approach, we assessed if contigs from short-read *de novo* assemblies of seven *T. urticae* strains (Table S1) bridged Sanger scaffolds. With pysam 0.14.1 (Li *et al.* 2009) and the BLASR 1.3.1 alignments of *de novo* short-read assemblies to Sanger scaffolds, we identified short-read assembled contigs across the seven strains for which at least 7.5 kb aligned to 75 kb segments (or the last 25% of the total scaffold length, whichever was smaller) at the ends of two different Sanger scaffolds. When this occurred, we joined the scaffolds into larger superscaffolds in an iterative manner (in a small number of cases, different contigs supported different joins; these were resolved based on best alignment support as described in the footnotes for Table S3). Finally, we resolved superscaffolds from the AWD-based joining and assembly-bridging approaches to produce three pseudochromosomes (pChr1-3). In constructing the final pseudochromosomes, we gave precedence to the assembly-bridging method as it made explicit predictions based on assembled sequences (Table S3, and see Results). Following pseudochromosome construction, gene coordinates from the Online Resource for Community Annotation of Eukaryotes (ORCAE) (Sterck *et al.* 2012) June 2016 *T. urticae* annotation were converted to pseudochromosome coordinates and checked, sorted and validated using GenomeTools 1.5.10 (Gremme *et al.* 2013).

### Bulked segregant analyses to detect responses to spirodiclofen

Our experimental design for spirodiclofen studies, in which eight paired selected/control populations were used, suggested a straightforward permutation approach for detection of significant responses to selection. For each pair of samples, we adapted methods from previous BSA studies of monogenic traits in *T. urticae* (Van Leeuwen *et al.* 2012; Demaeght *et al.* 2014; Bryon *et al.* 2017a) and allele frequencies as assessed for AWD calculations (see section “Identification of misassembled regions in Sanger scaffolds”) to calculate the genome-wide change in SR-VP allele frequencies between all selected and control pairs (the analysis was performed using pChr1-3). In BSA mapping studies, deviations in allele frequencies from zero occur by genetic drift (where they are expected to be uncorrelated between selected/control pairs), or in response to selection (where they should be correlated in location and direction of change) (Bryon *et al.* 2017b). To establish regions of correlated responses, we first averaged BSA scans from all eight pairs over the genome (this created the observed distribution as assessed with all replicate information). Then, we permuted the scan data for the eight replicates 10^4^ times; in each instance we calculated an analogous BSA average across the permuted eight replicates. To maintain linkage information for each permutation instance, we treated the concatenated genome (pChr1-3) as if it was circular and chose random start locations for each of the eight scans. From the 10^4^permutations, we then constructed a distribution of the absolute values of the maximal deviations in the averaged BSAs from zero (one data point per permutation; absolute values were taken as responses to drift and selection can be in the direction of either parent). We assigned as QTL those peaks where the observed allele frequency maxima were greater than the 95th percentile of values from the permutations (a false discovery rate, FDR, of 5%).

### BSA analyses to detect responses to tomato

BSA analyses and permutation-based detection of QTL for selection on tomato was performed as for spirodiclofen selections with one modification. As the five tomato/control populations were not paired, we generated all possible five-to-five combinations of tomato-selected and control populations. For each grouping, BSA scans were performed, and 10^4^ permutations were used to establish combination-specific values for detecting significant QTL (FDRs of 5%).

### Detection of QTL with the G’ method

As a complementary approach for QTL detection, we used the G’ method (Magwene *et al.* 2011) as implemented in QTLseqr 0.6.4. (Mansfeld and Grumet 2018). As input for QTLseqr, alignments from replicates from the respective spirodiclofen and tomato selections, and the respective control populations, were pooled for variant calling by GATK to form “high bulk” and “low bulk” groups, respectively; the resulting VCF file was converted into the table format using the GATK VariantsToTable tool. For quality control for variant selection, we followed the recommendations provided in the QTLseqr manual and vignette. We only included SNPs with a combined bulk read coverage of 400-500, a coverage of at least 200 for each bulk, and genotype quality scores of 99; additionally, SNPs were excluded from the analysis if they had a reference strain allele frequency of below 0.05 or above 0.95 in both high and low bulks, and fell outside the DeltaSNP filter threshold of 0.15 (spirodiclofen selection) and 0.10 (tomato selection). The DeltaSNP filter thresholds were empirically determined from analyses of the fits of the filtered data to null log G’ distributions as described in the QTLseqr vignette. Window sizes were set at 500 kb, and the genome-wide FDR for QTL intervals was set to 0.05.

### Analysis of candidate genes for responses to selection

Genetic differences between Lon-Inb and SR-VP in coding regions of candidate genes for response to spirodiclofen selection were annotated with SnpEff 4.2 (Cingolani *et al.* 2012). Predicted variants and their annotated effects on candidate genes were visually curated in Integrative Genomics Viewer 2.3 (Robinson *et al.* 2011; Thorvaldsdóttir *et al.* 2013) with alignments of Illumina reads for Lon-Inb and SR-VP, as well as with alignments of short-read *de novo* assemblies for these two strains.

To assess if nonsynonymous changes were unique to SR-VP, we examined alignments of genomic Illumina reads available from the seven additional strains reported in this study (Table S1), five strains or populations reported by Bryon *et al.* 2017a, strain EtoxR (Van Leeuwen *et al.*2012), strain HexR (Demaeght *et al.* 2014), and strain Montpellier (Grbić *et al.* 2011). We also tested for the presence of nonsynonymous variants unique to SR-VP in strain Harbin for which only RNA-seq data was available (Zhao *et al.* 2016). To do this, we generated alignments with the respective RNA-seq reads using the two-pass mode of STAR 2.5.2b (Dobin *et al.* 2013) with a maximum intron size of 20 kb. Visual assessment of one candidate region for response to spirodiclofen selection suggested extensive copy number variation. To quantify this, we assessed read coverage for both Lon-Inb and SR-VP throughout the region underlying the peak response and normalized it to the pseudochromosome-wide mean coverage as assessed from BAM files for Lon-Inb and SR-VP using pysam 0.9.1.4.

### Data availability

Sequence data has been deposited at the Sequence Read Archive (PRJNA498683). Supplemental figures and tables are available at FigShare. Other data, including variant loci and allele frequency information as a VCF file, BLASR-alignments of short-read assemblies to the Sanger reference assembly as BAM files, along with the respective input files for alignments, and the *T. urticae* pseudochromosome assembly, are available at the National Science Foundation supported CyVerse repository (public links for review are appended to this single document PDF submission; at acceptance, a permanent DOI for the data sets will be generated). Available strains will be distributed by the corresponding authors (permits may be required).

## RESULTS

### Phenotypic responses to pesticide and host plant selections

Our study used two inbred strains of *T. urticae*, SR-VP and Lon-Inb, which were derived from two previously characterized populations (Van Pottelberge *et al.* 2009; Grbić *et al.* 2011). As revealed by toxicity bioassays, the SR-VP strain maintained high-level resistance to spirodiclofen as found in its parental population (Van Pottelberge *et al.* 2009), while Lon-Inb was susceptible. The LC_50_ for Lon-Inb was a low 2.8 mg/L (95% confidence interval, 2.4 to 3.2 mg/L), while the LC_50_ value could not be calculated for SR-VP as resistance levels were too high (a reliable calculation would require concentrations higher than 5000 mg/L). Survival varied significantly at doses of both 2500 and 5000 mg/L spirodiclofen (each *p*-value < 0.0001 as identified by a generalized linear model with a binomial distribution; Figure 1A). In addition, reproductive performance of SR-VP on tomato was ~10-fold higher than for Lon-Inb, a significant difference (*p* < 0.0001 as identified by a general linear model, Figure 1B).

In experimental populations propagated for ~50 generations following a SR-VP × Lon-Inb cross, survival at both 2500 and 5000 mg/L of spirodiclofen was significantly higher for spirodiclofen-selected populations compared to their paired control populations, and survival differed between the two doses (generalized linear mixed model with a binomial distribution, *p*-values < 0.0001; Figure 1A). In response to growth on tomato, after ~50 generations tomato-selected mite populations had significantly higher performance on tomato as compared to the control populations maintained on bean (general linear mixed model, *p-*value = 0.0189 Figure 1B); however, in contrast to selection by spirodiclofen, the phenotypic difference was modest (compare Figure 1A to Figure 1B).

### Genomic responses to selection

To examine genomic responses to selection, we chose, based on large resistance ratios, eight pairs of spirodiclofen-selected and matching control populations for genomic analyses (Figure 1A). Additionally, we chose five tomato-selected populations with high performance on tomato (Figure 1B), and an additional control population that performed poorly on tomato (population C15; in total nine control populations were selected to inform responses to spirodiclofen, tomato plants, or both, Figure 1). To assess genomic responses to selection, we sequenced genomic DNA from these 22 populations, as well as the parental SR-VP and Lon-Inb strains.

At each of 694,308 high-quality SNP loci that distinguished SR-VP and Lon-Inb, we determined the frequency of the SR-VP allele in each of the 22 populations. A PCA with the resulting data revealed that all tomato-selected populations were distinct along principal component 1 (PC1) from control and spirodiclofen-selected populations; along PC2, spirodiclofen-selected populations clustered separately from control populations (Figure 2).

As controls and treatments were separated by a PCA, we examined allele frequencies of populations across the largest 44 scaffolds in the Sanger assembly, all of which are 185 kb or larger and collectively harbor ~95% of the assembly length (Grbić *et al.* 2011; Van Leeuwen *et al.* 2012). A sliding window analysis revealed that across all populations allele frequencies were broadly similar over much of the genome (Figure 4A). However, for several small regions, for example on Sanger scaffolds 5 and 16, fixation (or near fixation) of alleles from one parent was observed, potentially reflecting the purging of segregating deleterious variants (see Discussion).

**Figure 4.**
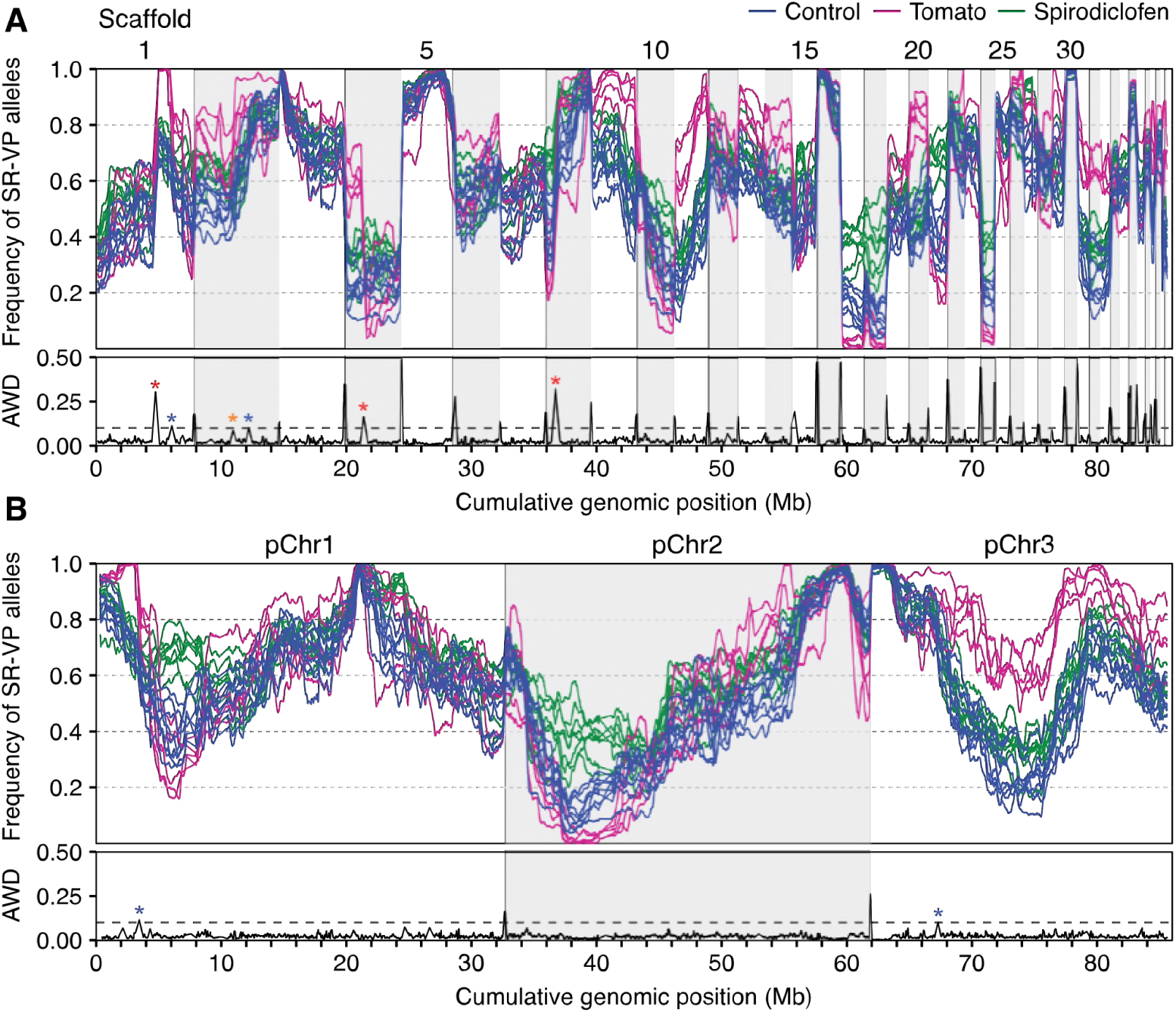
A pseudochromosome assembly resolves discontinuities in allele frequencies in experimental populations across the genome. (A and B) Frequency of SR-VP alleles in control (blue), spirodiclofen-selected (green), and tomato-selected (magenta) populations as assessed in a sliding window analysis (legend, top right). Allele frequencies are shown using the first 44 scaffolds of the Sanger assembly (A), or the consolidated pseudochromosome assembly (B). Concatenated Sanger scaffolds or pseudochromosomes are indicated by alternating white and gray shading, and are sorted by decreasing length. At the bottom of each panel, the respective average window distances (AWD) are shown as assessed with allele frequency data from all populations. In each panel, the dashed line represents an AWD value of 0.1 (a value suggestive of misassemblies, see Results section). Three AWD peaks well above the threshold correspond to obvious misassemblies (red asterisks on Sanger scaffolds 1, 4, and 8); the AWD peak on Sanger scaffold 2 denoted with an orange asterisk was identified as a misassembly previously (Bryon *et al.*, 2017a; see also Figure S3A). Two other peaks barely exceed the AWD value of 0.1 (on Sanger scaffolds 1 and 2, corresponding to pseudochromosomes 1 and 3 in the consolidated assembly, respectively; blue asterisks, A and B); these peaks were not supported as misassemblies in independent data sets (Figures S3 and S4), or in assemblies of *T. urticae* strains using short-read data.

Nevertheless, at some loci systematic deviations in allele frequencies were observed between control and selected populations, suggesting responses to selection. For example, in regions on Sanger scaffolds 17 and 21 all spirodiclofen-selected populations differed from control populations, and in regions on Sanger scaffolds 11 and 32, all tomato-selected populations differed from control populations (Figure 4A). However, some of these regions were present on small Sanger scaffolds, or were near the ends of larger ones. Therefore, comprehensive description of genomic responses to selection was not possible with the existing draft genome. Additionally, some larger Sanger scaffolds harbored marked discontinuities in allele frequencies, revealing putative misassemblies, as also reported in a previous study (Bryon *et al.* 2017a).

### Genome scaffolding with population genetic data

The sliding window analyses revealed the limitations of the existing draft Sanger assembly for our study, and also suggested a way to overcome them. We reasoned that similarities in population allele frequencies within and between Sanger scaffolds could be used to resolve misassembled regions, as well as determine relative scaffold positions in the genome. For example, as apparent from the allele frequency data in Figure 4A, Sanger scaffold 5 cannot possibly be adjacent to Sanger scaffolds 17 or 24 in the *T. urticae* genome, but it could plausibly be adjacent to Sanger scaffolds such as 16 or 22.

To identify misassembled regions, we constructed a metric, AWD, to assess the continuity of allele frequencies in experimental populations between non-overlapping genomic windows (Figure 3, and Materials and Methods). AWD values are expected to be small between adjacent, non-overlapping windows in correctly assembled genomic regions, as in our experimental populations major allele frequency changes occurred at Mb scales (i.e., see Sanger scaffold 3 in Figure 4A). Consistent with this expectation, and after correcting for a small set of windows that gave locally anomalous allele frequencies (a potential effect of incorrect variant predictions or structural variation between strains, Figure S1), AWD values between adjacent windows were close to zero in most genomic intervals (the mean value within Sanger scaffolds was 0.025; Figure 4A, bottom). Exceptions occurred at the junctions between Sanger scaffolds, which were concatenated by decreasing length as shown in Figure 4A and, except by chance, are not expected to be physically adjacent. As calculated between concatenated Sanger scaffolds, the mean AWD value was 0.243, with a range between ~0.10 to 0.50; these values establish a *de facto* expectation for the magnitude of an AWD value anticipated at the site of a misassembly.

Applying a conservative AWD threshold for detecting assembly errors (Figure S1), clear misassemblies were apparent within Sanger scaffolds 1, 4 and 8 (Figure 4A, red asterisks). In each case, these corresponded to misassemblies previously noted by Bryon *et al.* (2017a) in an unrelated BSA mapping study in *T. urticae*. Further, Bryon *et al.* (2017a) reported a misassembly on scaffold 2 at the location of a less dramatic but nonetheless elevated AWD value (orange asterisk in Figure 4A, and see Figure S1). For subsequent analyses, we treated these as candidate misassemblies, and broke the respective four Sanger scaffolds into subscaffolds (Table S2).

Linking together Sanger scaffolds with reciprocal minimal AWD values in terminal windows (Materials and Methods, and see Figure 3) generated three superscaffolds (hereafter referred to as “AWD-joined superscaffolds”). Each of the first 44 largest Sanger scaffolds was included in these superscaffolds except Sanger scaffolds 41 and 42, which were unplaced. Briefly, Sanger scaffold 41 was not polymorphic between SR-VP and Lon-Inb (Figure S2), and therefore could not be joined as the AWD method requires genetic differences. For Sanger scaffold 42, inspection of aligned Illumina reads revealed massive copy number variation (Figure S2), suggesting a complex misassembly; we therefore excluded it from subsequent analyses. Finally, of the Sanger scaffolds included in the AWD-joined superscaffolds, the orientations of those higher than 39 could not be determined (see Materials and Methods, Table S4).

### Sequence-based scaffolding of the Sanger assembly

As a complementary approach to condense Sanger scaffolds into superscaffolds, we also leveraged short-read *de novo* assemblies for SR-VP, Lon-Inb, and five additional *T. urticae* strains. As expected, these Illumina short-read assemblies were more fragmented than the Sanger assembly (Table S1). Given the potential for errors in short-read assemblies of many thousands of contigs, we did not attempt to systematically use these assemblies to identify errors internal to Sanger scaffolds. In a more limited analysis, however, we identified instances where contigs “bridged” two Sanger scaffolds, and joined all but three Sanger scaffolds (8.1, 21, and 25) into superscaffolds (hereafter termed “assembly-bridged superscaffolds”; Table S3). As compared to the AWD-joined superscaffolds, more assembly-bridged superscaffolds were produced (eight as opposed to three). In the assembly-bridged superscaffolds, Sanger scaffold 41, which was unplaced in the AWD-joined superscaffolds, was bridged to Sanger scaffold 36.

### Consolidation of assemblies to three pseudochromosomes

To produce a consolidated *T. urticae* pseudochromosome assembly, we resolved the AWD-joined and assembly-bridged superscaffolds (Figure 5). Where they could be compared, only a single discrepancy was observed. While in both sets of superscaffolds Sanger scaffolds 39 and 43 were together between the larger Sanger scaffolds 20 and 31 (Figure 5), the relative positions differed (Tables S3 and S4). The order from the assembly-bridged superscaffolds was selected for this pseudochromosome join, as well as for establishing the orientation of Sanger scaffolds 39 and higher (see Materials and Methods). Resolution of the two superscaffold assemblies resulted in three pseudochromosomes, pChr1-3, of lengths 32.7, 29.2 and 23.9 Mb, respectively (Figure 5). Further, short-read *de novo* assemblies were used to refine misassembly breakpoints in Sanger scaffolds 1, 2, 4, and 8 (Table S2).

**Figure 5.**
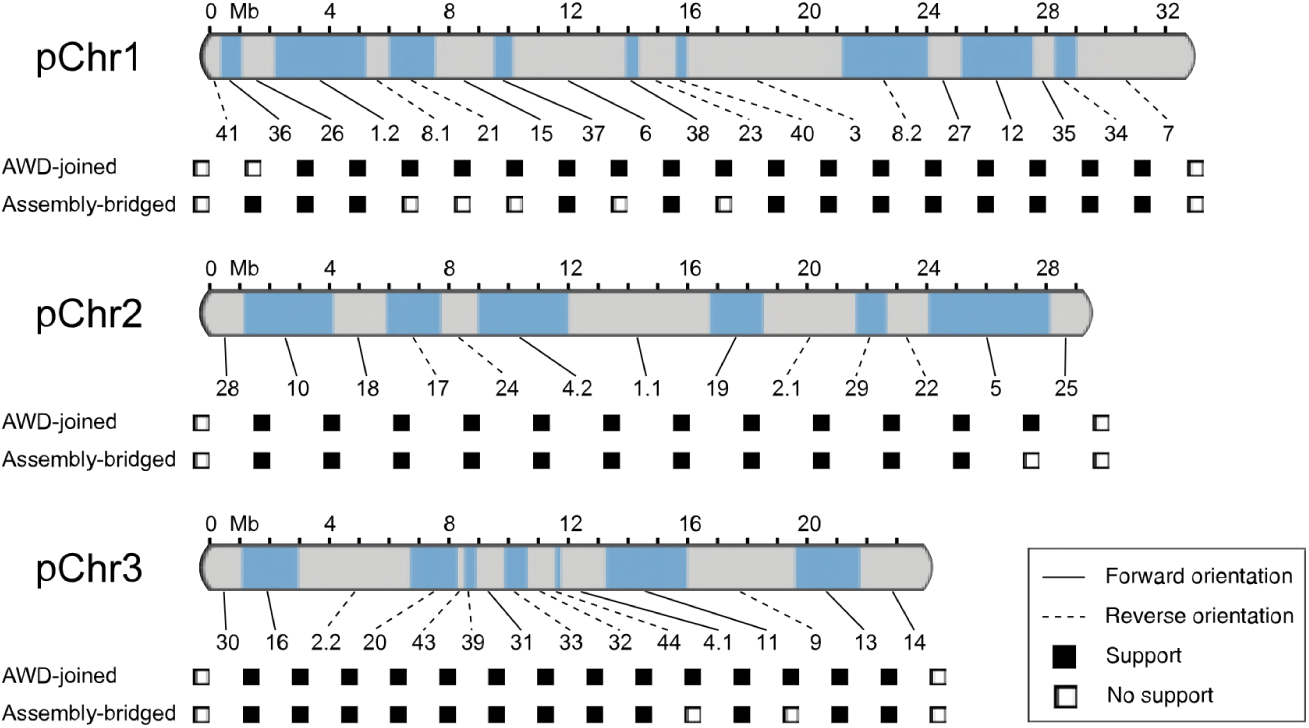
Pseudochromosomes constructed from AWD-joined and assembly-bridged superscaffolds. Composition of Sanger scaffolds in three pseudochromosomes (pChr1-3) with orientation indicated by solid or dashed lines (see legend; Sanger scaffolds are indicated by alternating blue and gray shading). The two sources of support for placing and orienting Sanger scaffolds – from the AWD-joining or assembly-bridging methods – are as indicated beneath each pseudochromosome. Contig numbers for respective *T. urticae* strains supporting assembly-bridging events are given in Table S3. For pChr3, AWD support is indicated as positive (filled squares) for the placement of Sanger scaffolds 39 and 43 between the larger Sanger scaffolds 20 and 31; however, the order and orientations of 39 and 43, as well as the orientation of other small Sanger scaffolds, was refined with short-read *de novo* assemblies.

### Validation of the pseudochromosome assembly

We performed several analyses to assess the validity of the three pseudochromosomes. First, we examined allele frequencies for the 22 control, spirodiclofen-, and tomato-selected populations along pChr1-3 (Figure 4B). As compared to Figure 4A, striking discontinuities in AWD values were no longer apparent. In two cases AWD values barely exceeded ~0.1, and in one of these cases, at 3.46 Mb on pChr1 (Figure 4B), the peak can be explained by a lack of genetic variation between SR-VP and Lon-Inb (a long shared haplotype between the strains in this region meant that the nearest adjacent windows available for AWD calculations were ~290 kb apart; hence, an elevated AWD value is expected). Further, on average, AWD values between all pairs of the ends of pChr1-3 were large (Table S5), consistent with correct chromosome end assignments (note that the short-read *de novo* assemblies also supported the ends assigned to pChr1-3, Figure 5 and Table S3, as no respective scaffolds from any *T. urticae* strains bridged any combination of the ends of pChr1-3).

Finally, we assessed the pseudochromosome assembly using two smaller population allele frequency data sets reported previously. First, we reanalyzed the data of Bryon *et al.* (2017a). In a sliding window analysis using the parameters employed in Figure 4, we found, as expected, marked discontinuities in population allele frequency data between unassembled Sanger scaffolds (Figure S3A). We also verified the misassemblies noted by Bryon *et al.* (2017a) on Sanger scaffolds 1, 2, 4 and 8, which were also observed in the current study (Figure 4A). When the analysis was repeated using the pseudochromosomes (Figure S3B), all major discontinuities disappeared, and only a handful of minor ones were apparent (e.g., in the middle of pChr3, potentially reflecting structural differences among strains, but see Discussion). An analysis of data from another study (Van Leeuwen *et al.* 2012) gave a similar result (Figure S4). Therefore, the pseudochromosome assembly resolved discontinuities in allele frequencies across the genome – an expectation of a correct assembly – in experimental data from two prior studies.

### Genomic regions underlying responses to spirodiclofen

With the three-pseudochromosome assembly, concerted increases in the frequency of SR-VP alleles were observed in spirodiclofen-treated as compared to control populations in several genomic regions (Figure 6A). To rule out an effect of genetic drift, we established a threshold for significant responses by permuting the data 10^4^ times (see Materials and Methods). At a FDR of 5%, two peaks on pChr1 (hereafter spiro-QTL 1, at 6.56 Mb, and spiro-QTL 2, at 24.13 Mb) and one peak on pChr2 (spiro-QTL 3 at 5.69 Mb) exceeded the significance threshold. In fact, the observed average allele frequency change of all eight pairwise contrasts (black line in Figure 6B) exceeded the maximum value from each of the 10^4^ permutations at each of spiro-QTL 1-3. Using the G’ approach as implemented in QTLseqr, these three peaks, and no others, were also identified as significant at a FDR of 5% (Figure S5).

**Figure 6.**
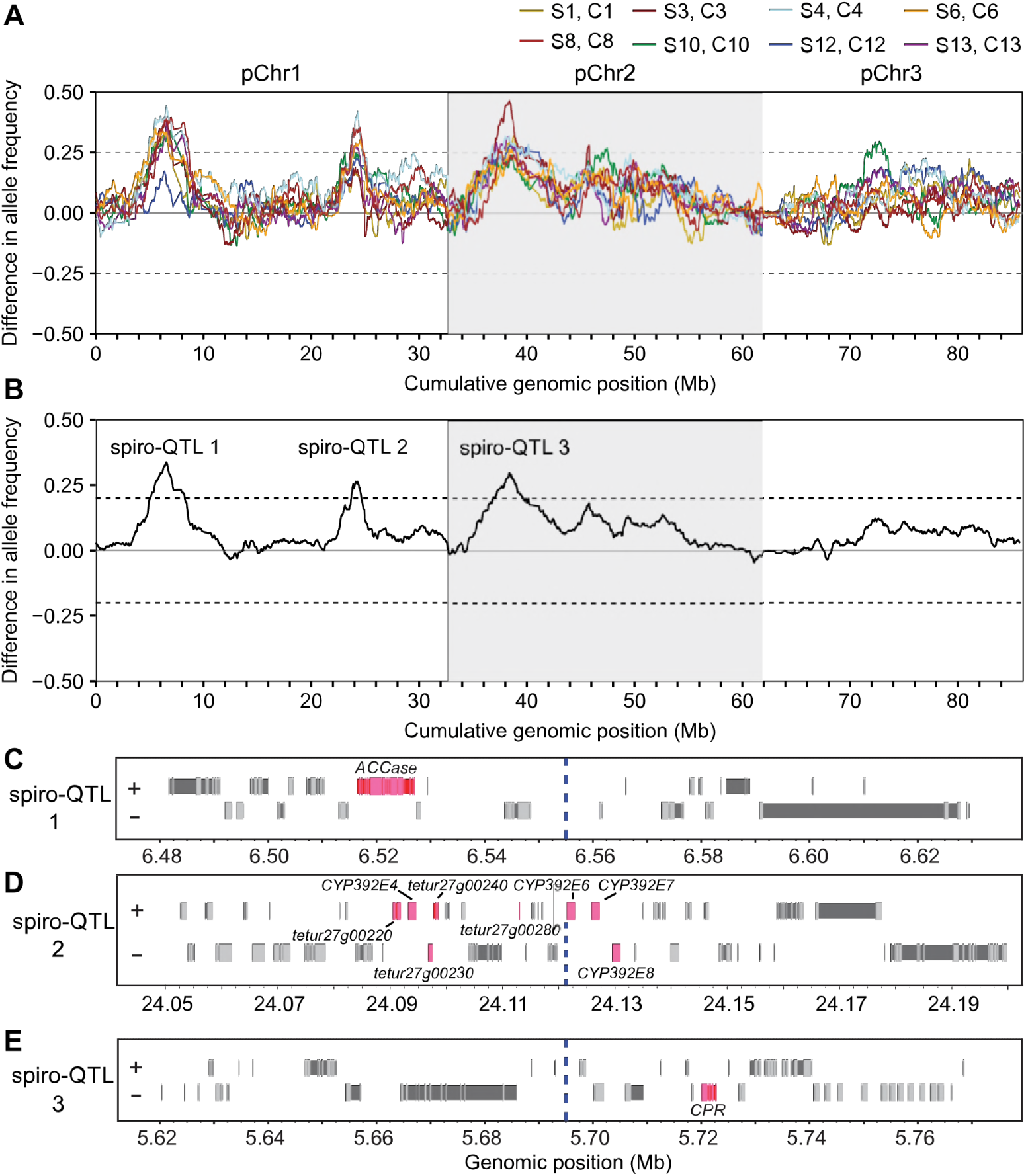
Genomic responses to spirodiclofen selection in long-term populations. (A) Genome-wide differences in SR-VP allele frequencies between pairs of eight spirodiclofen-selected populations and their matching control populations (legend, top right) as assessed with sliding windows. (B) Mean change in SR-VP allele frequency as assessed with all eight paired spirodiclofen/control replicates. Three QTL (spiro-QTL 1-3) exceed the 5% FDR threshold for detection of responses to selection as assessed by a permutation approach. (C-E) Gene models within the peak 150 kb windows for each of spiro-QTL 1-3. Coding exons and introns are represented by light gray and darker boxes, respectively (+ and – denote forward and reverse gene orientations). Candidate genes including *acetyl-CoA decarboxylase* (*ACCase*, *tetur21g02170*) (C), *CYP* genes (pseudogenes or fragments have “tetur” IDs) (D), and *NADPH cytochrome P450 reductase* (*CPR*, *tetur18g03390*) (E) are in pink. For panels C-E, coordinates are for the respective pseudochromosomes (Figure 5), and the vertical dashed lines denote the respective peaks shown in panel B.

To assess potential genes and variants responding to selection, we examined 150 kb genomic intervals centered on the QTL peak regions (Figure 6C-E). For spiro-QTL 1, no annotated detoxification genes were present; however, the peak was within 27.8 kb of *acetyl-CoA decarboxylase* (*ACCase*; *tetur21g02170*) (Figure 6C and Table S6), which encodes the target of spirodiclofen. As assessed with short-read alignments and *de novo* assemblies of Lon-Inb and SR-VP, for which contigs extended across the entire 7002 bp open reading frame of the large *ACCase* gene (contig numbers 849 and 1261 in the respective short-read assemblies), there were 37 single nucleotide differences. ACCase is highly conserved in eukaryotes, and consistent with purifying selection, 36 of these changes were synonymous. The single nonsynonymous change, an alanine to threonine change at position 1079 (A1079T), was unique to SR-VP compared to 16 other strains for which sequence data was available (see Materials and Methods).

In contrast, the peak region for spiro-QTL 2 was broader, forming a plateau of ~1 Mb in length (Figure 6B and Figure S6). The maximum change in allele frequencies at spiro-QTL 2 fell internal to a cluster of CYPs (Figure 6D). Four of these, *CYP392E4*, *CYP392E6*, *CYP392E7*, and *CYP392E8*, are intact in the Sanger reference sequence, one is an annotated pseudogene (*CYP392E5p*), and two additional annotations reflect apparent CYP fragments (*tetur27g00240* and *tetur27g00280*) (Table S7). Two other CYPs, *CYP392E9* and *CYP392E10*, which are present in a separate cluster, are located ~330 kb distal to the peak region within the ~1 Mb interval (Figure S6). For both SR-VP and Lon-Inb, no short-read *de novo* contigs spanned the CYP cluster at the peak (Figure 6D), and as revealed by read coverage depth the region harbors substantial structural variation between SR-VP and Lon-Inb (Figure S7). For example, in Lon-Inb, most of the CYPs are present in approximately two to seven copies relative to the Sanger reference sequence. In contrast, in SR-VP read coverage for most of the CYPs revealed that they are present as single copies.

Finally, although no *CYPs* or genes in other known detoxification families are present near spiro-QTL 3 (Figure 6E and Table S8), *NADPH cytochrome P450 reductase* (*CPR*, *tetur18g03390*), which is required for CYP activity, is located within ~25 kb of the sharp peak in allele frequency changes. As assessed with short-read alignments and *de novo* assemblies that spanned this locus (contig numbers 633 and 459 for Lon-Inb and SR-VP, respectively), 10 nucleotide changes between Lon-Inb and SR-VP were present in the 2001 bp coding region of *CPR.* Only one change in SR-VP impacted the coding sequence, an aspartic acid to tyrosine change at position 384 (D384Y). This variant was present in only one other strain, HexR.

### Genomic regions associated with selection by tomato

As opposed to the paired experimental design for selection studies with spirodiclofen, the five tomato-selected populations were unpaired to controls (Figure 1, and Materials and Methods). Therefore, we analyzed all possible groupings of the tomato-selected populations to five control populations. For each combination, we performed a QTL scan with the permutation approach as described for the spirodiclofen-selection analysis. In every combination, peaks that exceeded a 5% FDR for QTL detection were identified in a broad region near the middle of pChr3 (Table S9; changes in allele frequencies were always in the direction of the SR-VP parent). A representative result of one combination is shown in Figure 7. No other genomic regions were detected at the 5% FDR threshold in any combinations. The minimal region on pChr3 that responded to selection extended from ~7.48 to 17.02 Mb (Table S9). This region of 9.54 Mb comprises ~10% of the length of the genome. As assessed with the G’ method (FDR of 5%), this entire region was also strongly supported as a QTL interval (Figure S8). In addition, with the G’ approach, six other QTL intervals were identified, albeit with modest G’ values, of which the most notable was at ~2.5 Mb on pChr1. For all but one of these, allele frequency changes were in the direction of the SR-VP parent.

**Figure 7.**
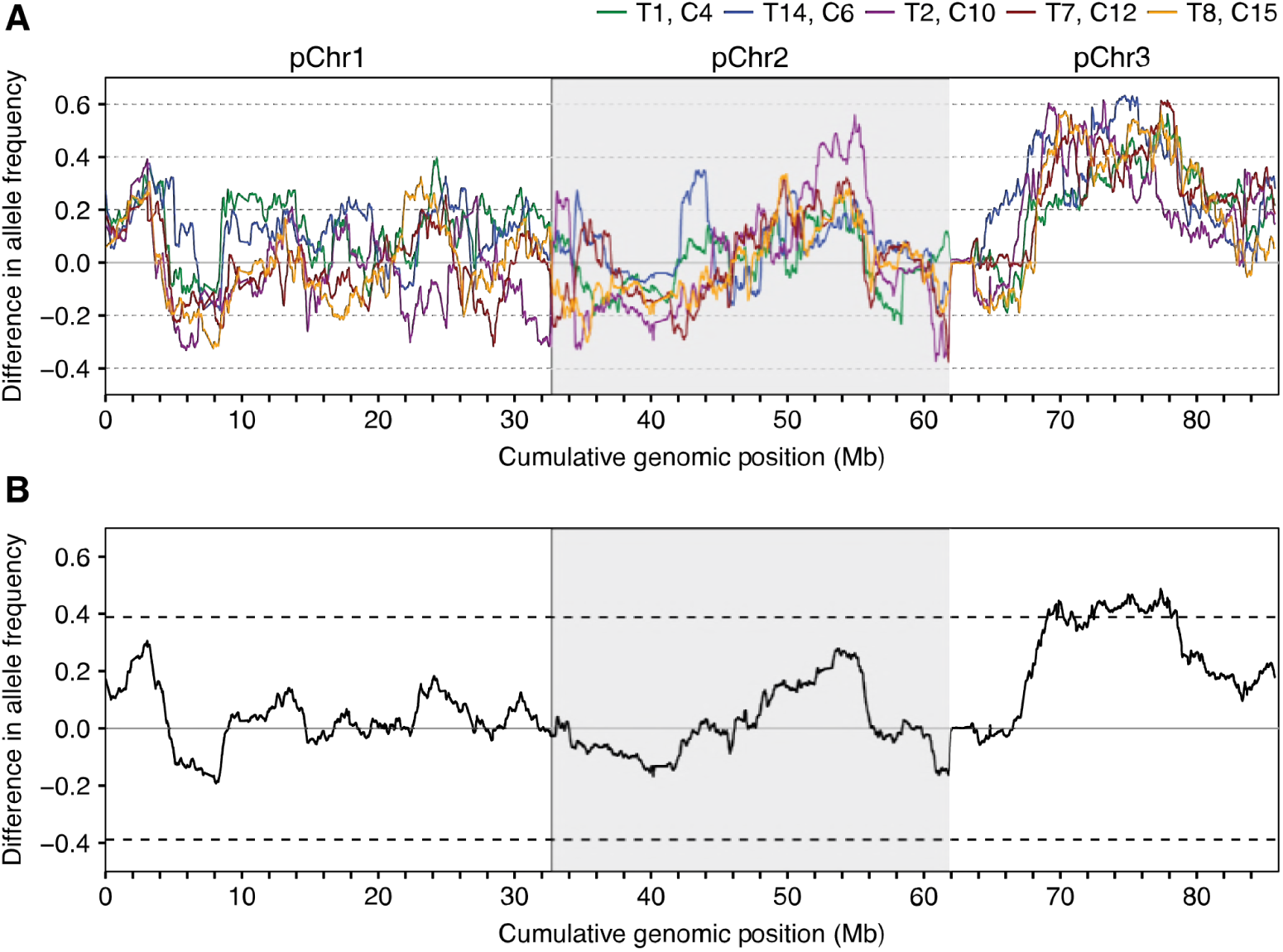
Genomic responses to selection on tomato plants in long-term populations. (A) Genome-wide differences in SR-VP allele frequencies between a representative pairing of five tomato-selected populations and five control populations (legend, top right) as assessed with sliding windows. (B) Mean change in SR-VP allele frequency as assessed with the five paired spirodiclofen/control replicates shown in panel A. Dashed lines denote a 5% FDR threshold for detection of responses to selection as assessed by a permutation approach.

## DISCUSSION

In earlier work, we developed and applied BSA methods to *T. urticae* populations to identify loci responsible for several monogenic traits (Van Leeuwen *et al.* 2012; Demaeght *et al.* 2014; Bryon *et al.* 2017a). In the current study, we extended these methods to reveal the quantitative basis of pesticide resistance and host plant adaptation by performing highly replicated, long-term experimental selections to spirodiclofen and tomato. As a first step, we resolved the draft Sanger assembly to three pseudochromosomes, a number matching the chromosome count reported from cytological studies (Helle and Bolland 1967). Notwithstanding new developments in long read sequencing technologies, assembling complete genomes of eukaryotes remains a challenge. The success of our population allele frequency approach to genome scaffolding results in part from the continuous nature of genome-wide allele frequency changes in populations. This contrasts with the more limited information that comes from individuals, e.g., from F2 mapping populations, where loci are either fixed for one of two alleles, or are heterozygous. Our genetic methods for curating and resolving draft assemblies should be applicable to other genome projects with species for which even a modest number of segregating populations can be generated.

Population allele frequency data from independent, future studies, potentially in combination with long read sequencing or optical mapping approaches (Shendure *et al.* 2017), will be important to identify and resolve any discrepancies in our refined *T. urticae* genome assembly. However, the pChr1-3 assembly resolved ambiguities in two prior *T. urticae* BSA datasets, and is therefore sufficient to assess genome-wide responses to selection with no (or little) uncertainty arising from assembly errors. As revealed from genome-wide allele frequency data, selection in all experimental populations was apparent, with the complete fixation of parental SR-VP haplotypes at several chromosomal locations (shifts toward alleles from the Lon-Inb parent were also apparent, but less extreme). These instances of fixation likely reflected the purging of deleterious alleles contributed by one or the other inbred parent, although our data do not exclude phenomena such as segregation distortion. Although these fixation events likely precluded our ability to detect responses to selection by spirodiclofen and tomato in these regions, they comprised only a tiny fraction of the genome.

In response to selection by spirodiclofen, three genomic regions responded significantly. In two of the three cases (spiro-QTL 1 and 3), sharp peaks in BSA scans were observed. This pattern is reminiscent of the ultra-high resolution BSA genetic mapping of monogenic loci observed in earlier *T. urticae* studies, in which the peaks of response were either within or only a few tens of kb from causal genes (Van Leeuwen *et al.* 2012; Demaeght *et al.* 2014; Bryon *et al.* 2017a). For each spiro-QTL, striking candidate genes were located within or immediately adjacent to replicate-averaged BSA peaks; these are discussed as follows, with the caveat that we cannot rule out the involvement of other genes within the QTL intervals.

The peak for spiro-QTL 1 was located nearby *ACCase*, the putative molecular target of spirodiclofen, raising the possibility of target-site resistance. This result was unexpected as an earlier study that examined coding sequences and expression of *ACCase* in spirodiclofen resistant *T. urticae* strains found no evidence for target-site resistance (Van Pottelberge *et al.* 2009). As opposed to the sharp peak of response at *ACCase*, the response at spiro-QTL 2 was broader, encompassing a region of ~600 kb. This genomic interval harbors two clusters of *CYPs* of the proliferated *T. urticae* CYP392 family (clan 2; genes *CYP392E4*, *-E6*, *-E7*, *-E8*, *-E9*, and *-E10*) (Grbić *et al.* 2011). Consistent with a role for CYPs in spirodiclofen resistance, treatment of the parental population from which SR-VP was derived with piperonyl butoxide (a CYP inhibitor) partially restored spirodiclofen sensitivity (Van Pottelberge *et al.* 2009). Further, *CYP392E7* and *CYP392E10* were also shown to be constitutively overexpressed and inducible by spirodiclofen in resistant *T. urticae* strains, and CYP392E10 was shown to metabolize spirodiclofen by hydroxylation in a heterologous assay (Van Pottelberge *et al.* 2009; Demaeght *et al.* 2013).

A prominent role for CYP-mediated detoxification of spirodiclofen is further supported by the finding that *CPR* is located at spiro-QTL 3. Previous studies demonstrated that the single copy of CPR in *T. urticae* is functional as a redox partner for CYPs (Demaeght *et al.* 2013; Riga *et al.* 2015). Nevertheless, while variation in CYPs has been associated with xenobiotic resistance in many animals, including spider mites (Van Leeuwen and Dermauw 2016), the finding that *CPR* is located at spiro-QTL 3 was not anticipated *a priori*. In particular, functional variation in CPR would be expected to affect multiple (or potentially all) CYPs, including those involved in primary metabolism and development. Therefore, allelic variants affecting CPR activity might be expected to be very deleterious. Some support for this conjecture comes from the control and tomato-adapted populations, which were not exposed to spirodiclofen, and where the SR-VP haplotype of *CPR* went to near extinction. However, purifying selection acting at a linked locus (or loci) cannot be excluded.

Despite the identification of loci and candidate genes for spirodiclofen resistance, the nature of underlying causal alleles, and how they interact to result in high-level resistance, remains unclear. Notwithstanding its widespread use, resistance to spirodiclofen is still comparatively rare. In this context, it is noteworthy that in SR-VP a single nonsynonymous change was present in *ACCase* that was absent from all other *T. urticae* strains that we examined (A1079T). A similar finding was observed for the D384Y variant in CPR, although the change was present in one other strain, HexR. Whether HexR is resistant to spirodiclofen is unknown, but CYP activity has been associated with resistance to many compounds, and HexR is a documented multi-pesticide resistant strain (Demaeght *et al.* 2014). How the amino acid changes in ACCase and CPR might impact resistance is not clear. For instance, the A1079T change in ACCase is outside the carboxyl-transferase domain that has been suggested to interact with keto-enol insecticides such as spirodiclofen (Lümmen *et al.* 2014). Hence, if A1079T contributes to target-site insensitivity, it must presumably do so through an allosteric mechanism. Assessing if the SR-VP variants in *ACCase* and *CPR* impact resistance will require further study, as will establishing whether CYP392E7, CYP392E10, or other CYP392 family members in the spiro-QTL 2 region metabolize spirodiclofen or its enol derivative *in vivo*. It should also be noted that substantial copy number variation for CYP392 family members was present between Lon-Inb and SR-VP (Figure S7), and short-read *de novo* assemblies for neither SR-VP nor Lon-Inb spanned the CYP cluster including *CYP392E4, −E6, −E7,* and *−E8*. Whether additional CYP genes are present in this region in Lon-Inb or SR-VP warrants further investigation. Regardless, our findings add to a growing body of evidence that copy number variation may play a prominent role in xenobiotic resistance in arthropods (Kwon *et al.* 2010; Zimmer *et al.* 2018; Weetman *et al.* 2018).

Finally, allele frequency shifts at each of the three spiro-QTL regions were comparatively modest (~25-35%). In only one case (spiro-QTL 2) did the SR-VP haplotype go to near fixation, although this haplotype was at a comparatively high frequency in control populations as well. The mechanism by which moderate changes in allele frequencies at a small number of loci can confer comparatively high resistance levels will require further study. Despite the small size of spider mites (~0.6 mm), single crosses can be performed, and recent work succeeded in constructing near-isogenic *T. urticae* lines to functionally validate the contribution of several target-site mutations to resistance phenotypes (Bajda *et al.* 2017; Riga *et al.* 2017). Fine-mapping and near-isogenic line construction is therefore possible, and will be important as a tool for understanding components of genetic architecture, like dominance and epistasis, that while important for quantitative resistance, are not straightforward to test in population-level selection experiments with pooled individuals.

While a primary focus of our study was on resistance to spirodiclofen, SR-VP’s performance on tomato relative to Lon-Inb was ~10-fold higher, which allowed us to test if genomic responses to selection by a pesticide and host plant are similar. Although far less than that observed for spirodiclofen resistance, the order of magnitude difference in tomato performance is comparatively large as assessed against prior studies of *T. urticae* strains on various host plants (Agrawal 2000; Wybouw *et al.* 2015). Even though our experimental design to assess selection by tomato was less powerful compared to that for spirodiclofen (paired samples were not used, and there were fewer replicates), as assessed by a PCA, tomato-selected populations were more strongly differentiated from control populations than were spirodiclofen-selected populations (i.e., genome-wide allele frequencies for spirodiclofen-selected populations more closely mirrored control populations than did tomato-selected ones, excluding the spiro-QTL 1-3 intervals). The genetic differentiation of tomato-selected samples, especially on pChr3, aided in genome reconstruction with the AWD method, and further a portion of pChr3 was significant for response to selection by tomato as assessed by permutations and the G’ method. The latter method also suggested additional responses to selection on all three pseudochromosomes; however, these results should be approached with caution as these regions did not reach the threshold for detection as established by permutation, and the G’ method was developed for and previously applied for QTL detection with simpler genetic designs (Magwene *et al.* 2011; Mansfeld and Grumet 2018).

Although additional work is required to validate the genetic basis of response to selection by tomato in the SR-VP × Lon-Inb cross, our findings suggest that it is likely highly polygenic. Further, the large candidate region for response to tomato on pChr3 was not detected as a significant QTL interval for selection by spirodiclofen, and the three QTL regions for response to spirodiclofen did not respond (or respond strongly) to selection by tomato. These findings suggest different genetic architectures. Our interpretation of a highly polygenic response to a host plant shift is consistent with several studies for host adaptation in plant-feeding insects (Jones 1998; Oppenheim *et al.* 2012). This may reflect the challenge that herbivores face in overcoming the many defensive and nutritional barriers plants have evolved to deter them (Strong *et al.* 1984; Howe and Jander 2008).

### Concluding remarks

Arthropod herbivores can adapt to novel host plants and pesticides. Deciphering the polygenic basis of these adaptation processes is important to maximize the effectiveness of modern integrated pest management strategies. We show that polygenic pesticide resistance and host plant use can be readily mapped in *T. urticae* using population-level selections, and that they can involve different genetic architectures. Further, genomic regions harboring small sets of candidate genes can be identified, an effect of recombination that accrues over multiple generations in populations of this herbivore. Although recent transcriptomic studies in arthropods have associated many gene families with xenobiotic resistance (Van Leeuwen and Dermauw 2016), our results with spirodiclofen nonetheless suggest a predominant role for genetic variation affecting a single major detoxification gene family (CYPs), potentially in combination with target-site resistance. In contrast, tomato adaptation was associated with the selection of a large genomic region, raising the possibility of more diverse underlying mechanisms. Studies with additional pesticide resistant and host-adapted strains will be important to extend our findings, as well as to establish their generality. Our work shows that such future studies are imminently possible in *T. urticae*, and in this context, our resolution of the genome assembly to pseudochromosomes should be invaluable.

## Acknowledgements

We thank Jon Seger and Fred Adler for helpful suggestions for detecting QTL with population allele frequency data, Ludek Tikovsky and Harold Lemereis for their assistance in plant rearing and maintenance of greenhouse rooms, Peter Demaeght for generation of the SR-VP inbred strain and René Feyereisen for helpful comments on the draft manuscript. This work was supported by the USA National Science Foundation (award 1457346 to R.M.C.) and the Research Foundation - Flanders (FWO, Belgium; grant G009312N to T.V.L. and grant G053815N to T.V.L. and W.D.). This project has received funding form the European Research Council (ERC) under the European Union’s Horizon 2020 research and innovation programme (grant agreement No 772026). N.W. was supported by a Marie Sklodowska-Curie Action (MSCA) Individual Fellowship (658795-DOGMITE) and a Research Foundation - Flanders (FWO) fellowship (12T9818N) throughout this project. R.G. was funded by National Institutes of Health Genetics Training Grant T32GM007464. W.D. is a postdoctoral fellow of the Research Foundation - Flanders (FWO). Research reported in this publication utilized the High-Throughput Genomics and Bioinformatic Analysis Shared Resource at Huntsman Cancer Institute at the University of Utah and was supported by the National Cancer Institute of the National Institutes of Health under Award Number P30CA042014. The content is solely the responsibility of the authors and does not necessarily represent the official views of the funding agencies.

*Large data sets to be hosted at the US National Science Foundation funded CyVerse site (public links are provided for access while the manuscript is under consideration; at acceptance, a DOI will be provided for the collective data files, and will be provided in the “Data Availability” statement)*.

## FILES

**Tomato_Spirodiclofen_Joint.3.6-0-g89b7209.vcf.gz**, a VCF file with genotypic data for parental strains and experimental populations. Link:

https://de.cyverse.org/dl/d/11CC290E-A900-4DF9-A233-1F71F3D2C693/Tomato_Spirodiclofen_Joint.3.6-0-g89b7209.vcf.gz

Heber.bam, a BLASR alignment of short-read assembled scaffolds from strain Heber to the *T. urticae* Sanger reference assembly. Link:

https://de.cyverse.org/dl/d/06735A0C-0B53-4C84-BBC8-B25AE38A617B/Heber.bam

**Heber.fasta.gz**, the input file used to generate file Heber.bam. Link:

https://de.cyverse.org/dl/d/5AFA43A1-3E0A-4A00-818F-E024F1BEE619/Heber.fasta.gz

Lon-Inb.bam, a BLASR alignment of short-read assembled scaffolds from strain Lon-Inb to the *T. urticae* Sanger reference assembly. Link:

https://de.cyverse.org/dl/d/E5A81F52-2CCF-4490-B462-FA032FDF7910/Lon-Inb.bam

**Lon-Inb.fasta.gz**, the input file used to generate file Lon-Inb.bam. Link:

https://de.cyverse.org/dl/d/369C3498-911F-47A0-8A27-C3D4477CF79E/Lon-Inb.fasta.gz

Parrott.bam, a BLASR alignment of short-read assembled scaffolds from strain Parrott to the *T. urticae* Sanger reference assembly. Link:

https://de.cyverse.org/dl/d/1FB6B2DB-CCFE-46C1-A9C7-B61E4DEB5CEF/Parrott.bam

**Parrott.fasta.gz**, the input file used to generate file Parrott.bam. Link:

https://de.cyverse.org/dl/d/6FA958FB-54E3-4F5B-9328-83DBE97F309C/Parrott.fasta.gz

RS.bam, a BLASR alignment of short-read assembled scaffolds from strain RS to the *T. urticae* Sanger reference assembly. Link:

https://de.cyverse.org/dl/d/07C5AA0E-9684-4F44-A9D0-5699E5373950/RS.bam

**RS.fasta.gz**, the input file used to generate file RS.bam. Link:

https://de.cyverse.org/dl/d/363ADA78-BB75-478B-AEC4-A5C86EB0903D/RS.fasta.gz

SR-VP.bam, a BLASR alignment of short-read assembled scaffolds from strain SR-VP to the *T. urticae* Sanger reference assembly. Link:

https://de.cyverse.org/dl/d/9506FBDE-5874-4822-8715-CBBA420F89DB/SR-VP.bam

**SR-VP.fasta.gz**, the input file used to generate file SR-VP.bam. Link:

https://de.cyverse.org/dl/d/78140292-BC67-459C-B5A9-FCCC84FC1E43/SR-VP.fasta.gz

**ShCo.bam**, a BLASR alignment of short-read assembled scaffolds from strain ShCo to the *T. urticae* Sanger reference assembly. Link:

https://de.cyverse.org/dl/d/71BB8C0E-12B5-4EDE-8F7C-D0933A11D3EF/ShCo.bam

**ShCo.fasta.gz**, the input file used to generate file ShCo.bam. Link:

https://de.cyverse.org/dl/d/116D52AC-1463-489C-A2EE-BD4FBBB800F6/ShCo.fasta.gz

**WG-Del.bam**, a BLASR alignment of short-read assembled scaffolds from strain WG-Del to the *T. urticae* Sanger reference assembly. Link:

https://de.cyverse.org/dl/d/4F50B8C5-8440-43EC-9FCE-7A2DBBE5DD1F/WG-Del.bam

**WG-Del.fasta.gz**, the input file used to generate file WG-Del.bam. Link:

https://de.cyverse.org/dl/d/A546E0BC-B0B6-48C5-BFB0-7D48954A4ACF/WG-Del.fasta.gz

**Pseudochromosome.fasta.gz**, the *T. urticae* pseudochromosome 1-3 sequences.

https://de.cyverse.org/dl/d/0FAB6310-135C-4F33-96F3-FEE77D5A3F06/Tetranychus_urticae.Pseudochromosome.fasta.gz

## REFERENCES

Agrawal A. A., 2000 Host-range evolution: adaptation and trade-offs in fitness of mites on alternative hosts. Ecology 81: 500–508.

Agrawal A. A., F. Vala, and M. W. Sabelis, 2002 Induction of Preference and Performance after Acclimation to Novel Hosts in a Phytophagous Spider Mite: Adaptive Plasticity? The American Naturalist 159: 553–565. https://doi.org/10.1086/339463

Alexandre H., S. Ponsard, D. Bourguet, R. Vitalis, P. Audiot, et al., 2013 When history repeats itself: exploring the genetic architecture of host-plant adaptation in two closely related lepidopteran species. PLoS ONE 8: e69211. https://doi.org/10.1371/journal.pone.0069211

Bajda S., W. Dermauw, R. Panteleri, N. Sugimoto, V. Douris, et al., 2017 A mutation in the PSST homologue of complex I (NADH:ubiquinone oxidoreductase) from Tetranychus urticae is associated with resistance to METI acaricides. Insect Biochemistry and Molecular Biology 80: 79–90. https://doi.org/10.1016/j.ibmb.2016.11.010

Bansal R., M. A. R. Mian, O. Mittapalli, and A. P. Michel, 2014 RNA-Seq reveals a xenobiotic stress response in the soybean aphid, Aphis glycines, when fed aphid-resistant soybean. BMC genomics 15: 972.

Bass C., C. T. Zimmer, J. M. Riveron, C. S. Wilding, C. S. Wondji, et al., 2013 Gene amplification and microsatellite polymorphism underlie a recent insect host shift. Proceedings of the National Academy of Sciences 110: 19460–19465. https://doi.org/10.1073/pnas.1314122110

Bretschneider T., R. Fisher, and R. Nauen, 2007 Inhibitors of lipid synthesis (acetyl-CoA-carboxylase inhibitors), pp. 909–925 in Modern crop protection compounds,.

Bryon A., A. H. Kurlovs, W. Dermauw, R. Greenhalgh, M. Riga, et al., 2017a Disruption of a horizontally transferred phytoene desaturase abolishes carotenoid accumulation and diapause in Tetranychus urticae. Proceedings of the National Academy of Sciences 114: E5871–E5880. https://doi.org/10.1073/pnas.1706865114

Bryon A., A. H. Kurlovs, T. Van Leeuwen, and R. M. Clark, 2017b A molecular-genetic understanding of diapause in spider mites: current knowledge and future directions: Molecular genetics of mite diapause. Physiological Entomology 42: 211–224. https://doi.org/10.1111/phen.12201

Ceccatti J. S., 2009 Insecticide resistance, economic entomology, and the evolutionary synthesis, 1914–1951. Trans. Am. Philos. Soc 1–21.

Chaisson M. J., and G. Tesler, 2012 Mapping single molecule sequencing reads using basic local alignment with successive refinement (BLASR): application and theory. BMC Bioinformatics 13: 238. https://doi.org/10.1186/1471-2105-13-238

Cingolani P., A. Platts, L. L. Wang, M. Coon, T. Nguyen, et al., 2012 A program for annotating and predicting the effects of single nucleotide polymorphisms, SnpEff: SNPs in the genome of Drosophila melanogaster strain w^1118^; iso-2; iso-3. Fly 6: 80–92. https://doi.org/10.4161/fly.19695

Coates B. S., and B. D. Siegfried, 2015 Linkage of an ABCC transporter to a single QTL that controls Ostrinia nubilalis larval resistance to the Bacillus thuringiensis Cry1Fa toxin. Insect Biochemistry and Molecular Biology 63: 86–96. https://doi.org/10.1016/j.ibmb.2015.06.003

Coates B. S., A. P. Alves, H. Wang, X. Zhou, T. Nowatzki, et al., 2016 Quantitative trait locus mapping and functional genomics of an organophosphate resistance trait in the western corn rootworm, Diabrotica virgifera virgifera. Insect Mol Biol 25: 1–15. https://doi.org/10.1111/imb.12194

Demaeght P., W. Dermauw, D. Tsakireli, J. Khajehali, R. Nauen, et al., 2013 Molecular analysis of resistance to acaricidal spirocyclic tetronic acids in Tetranychus urticae: CYP392E10 metabolizes spirodiclofen, but not its corresponding enol. Insect Biochemistry and Molecular Biology 43: 544–554. https://doi.org/10.1016/j.ibmb.2013.03.007

Demaeght P., E. J. Osborne, J. Odman-Naresh, M. Grbić, R. Nauen, et al., 2014 High resolution genetic mapping uncovers chitin synthase-1 as the target-site of the structurally diverse mite growth inhibitors clofentezine, hexythiazox and etoxazole in Tetranychus urticae. Insect Biochemistry and Molecular Biology 51: 52–61. https://doi.org/10.1016/j.ibmb.2014.05.004

Dermauw W., N. Wybouw, S. Rombauts, B. Menten, J. Vontas, et al., 2013 A link between host plant adaptation and pesticide resistance in the polyphagous spider mite Tetranychus urticae. Proceedings of the National Academy of Sciences 110: E113–E122. https://doi.org/10.1073/pnas.1213214110

Dermauw W., A. Pym, C. Bass, T. Van Leeuwen, and R. Feyereisen, 2018 Does host plant adaptation lead to pesticide resistance in generalist herbivores? Current Opinion in Insect Science 26: 25–33. https://doi.org/10.1016/j.cois.2018.01.001

Díaz-Riquelme J., V. Zhurov, C. Rioja, I. Pérez-Moreno, R. Torres-Pérez, et al., 2016 Comparative genome-wide transcriptome analysis of Vitis vinifera responses to adapted and non-adapted strains of two-spotted spider mite, Tetranyhus urticae. BMC Genomics 17: 74. https://doi.org/10.1186/s12864-016-2401-3

Dobin A., C. A. Davis, F. Schlesinger, J. Drenkow, C. Zaleski, et al., 2013 STAR: ultrafast universal RNA-seq aligner. Bioinformatics 29: 15–21. https://doi.org/10.1093/bioinformatics/bts635

Dobzhansky T., 1937 Genetics and the Origin of Species. Columbia University Press, New York.

Douris V., D. Steinbach, R. Panteleri, I. Livadaras, J. A. Pickett, et al., 2016 Resistance mutation conserved between insects and mites unravels the benzoylurea insecticide mode of action on chitin biosynthesis. Proceedings of the National Academy of Sciences 113: 14692–14697. https://doi.org/10.1073/pnas.1618258113

Feyereisen R., W. Dermauw, and T. Van Leeuwen, 2015 Genotype to phenotype, the molecular and physiological dimensions of resistance in arthropods. Pestic Biochem Physiol 121: 61–77. https://doi.org/10.1016/j.pestbp.2015.01.004

ffrench-Constant R. H., P. J. Daborn, and G. Le Goff, 2004 The genetics and genomics of insecticide resistance. Trends Genet. 20: 163–170. https://doi.org/10.1016/j.tig.2004.01.003

Fry J. D., 1989 Evolutionary adaptation to host plants in a laboratory population of the phytophagous mite Tetranychus urticae Koch. Oecologia 81: 559–565.

Gahan L. J., 2001 Identification of a Gene Associated with Bt Resistance in Heliothis virescens. Science 293: 857–860. https://doi.org/10.1126/science.1060949

Gould F., 1979 Rapid host range evolution in a population of the phytophagous mite Tetranychus urticae Koch. Evolution 33: 791. https://doi.org/10.2307/2407646

Grbić M., A. Khila, K.-Z. Lee, A. Bjelica, V. Grbić, et al., 2007 Mity model:Tetranychus urticae, a candidate for chelicerate model organism. BioEssays 29: 489–496. https://doi.org/10.1002/bies.20564

Grbić M., T. Van Leeuwen, R. M. Clark, S. Rombauts, P. Rouzé, et al., 2011 The genome of Tetranychus urticae reveals herbivorous pest adaptations. Nature 479: 487–492. https://doi.org/10.1038/nature10640

Gremme G., S. Steinbiss, and S. Kurtz, 2013 GenomeTools: A Comprehensive Software Library for Efficient Processing of Structured Genome Annotations. IEEE/ACM Transactions on Computational Biology and Bioinformatics 10: 645–656. https://doi.org/10.1109/TCBB.2013.68

Hardy N. B., D. A. Peterson, L. Ross, and J. A. Rosenheim, 2018 Does a plant-eating insect’s diet govern the evolution of insecticide resistance? Comparative tests of the pre-adaptation hypothesis. Evolutionary Applications 11: 739–747. https://doi.org/10.1111/eva.12579

Hawkins N. J., C. Bass, A. Dixon, and P. Neve, 2018 The evolutionary origins of pesticide resistance. Biological Reviews of the Cambridge Philosophical Society. https://doi.org/10.1111/brv.12440 [Epub ahead of print]

Helle W., and H. R. Bolland, 1967 Karyotypes and sex-determination in spider mites (Tetranychidae). Genetica 38: 43–53.

Hemingway J., N. J. Hawkes, L. McCarroll, and H. Ranson, 2004 The molecular basis of insecticide resistance in mosquitoes. Insect Biochemistry and Molecular Biology 34: 653–665. https://doi.org/10.1016/j.ibmb.2004.03.018

Henniges-Janssen K., A. Reineke, D. G. Heckel, and A. T. Groot, 2011 Complex inheritance of larval adaptation in Plutella xylostella to a novel host plant. Heredity (Edinb) 107: 421–432. https://doi.org/10.1038/hdy.2011.27

Howe G. A., and G. Jander, 2008 Plant immunity to insect herbivores. Annual Review of Plant Biology 59: 41–66. https://doi.org/10.1146/annurev.arplant.59.032607.092825

Jaquiéry J., S. Stoeckel, P. Nouhaud, L. Mieuzet, F. Mahéo, et al., 2012 Genome scans reveal candidate regions involved in the adaptation to host plant in the pea aphid complex. Mol. Ecol. 21: 5251–5264. https://doi.org/10.1111/mec.12048

Jones C. D., 1998 The genetic basis of Drosophila sechellia’s resistance to a host plant toxin. Genetics 149: 1899–1908.

Kwon D. H., J. M. Clark, and S. H. Lee, 2010 Extensive gene duplication of acetylcholinesterase associated with organophosphate resistance in the two-spotted spider mite. Insect Molecular Biology 19: 195–204. https://doi.org/10.1111/j.1365-2583.2009.00958.x

Li X., M. A. Schuler, and M. R. Berenbaum, 2007 Molecular Mechanisms of Metabolic Resistance to Synthetic and Natural Xenobiotics. Annual Review of Entomology 52: 231–253. https://doi.org/10.1146/annurev.ento.51.110104.151104

Li H., B. Handsaker, A. Wysoker, T. Fennell, J. Ruan, et al., 2009 The Sequence Alignment/Map format and SAMtools. Bioinformatics 25: 2078–2079. https://doi.org/10.1093/bioinformatics/btp352

Li H., 2013 Aligning sequence reads, clone sequences and assembly contigs with BWA-MEM. arXiv:1303.3997 [q-bio].

Lümmen P., J. Khajehali, K. Luther, and T. Van Leeuwen, 2014 The cyclic keto-enol insecticide spirotetramat inhibits insect and spider mite acetyl-CoA carboxylases by interfering with the carboxyltransferase partial reaction. Insect Biochemistry and Molecular Biology 55: 1–8. https://doi.org/10.1016/j.ibmb.2014.09.010

Magalhães S., J. Fayard, A. Janssen, D. Carbonell, and I. Olivieri, 2007 Adaptation in a spider mite population after long-term evolution on a single host plant. J. Evol. Biol. 20: 2016–2027. https://doi.org/10.1111/j.1420-9101.2007.01365.x

Magalhães S., E. Blanchet, M. Egas, and I. Olivieri, 2009 Are adaptation costs necessary to build up a local adaptation pattern? BMC Evol. Biol. 9: 182. https://doi.org/10.1186/1471-2148-9-182

Magwene P. M., J. H. Willis, and J. K. Kelly, 2011 The statistics of bulk segregant analysis using next generation sequencing. PLoS Computational Biology 7: e1002255. https://doi.org/10.1371/journal.pcbi.1002255

Mansfeld B. N., and R. Grumet, 2018 QTLseqr: An R package for bulk segregant analysis with next-generation sequencing. The Plant Genome 11: 0. https://doi.org/10.3835/plantgenome2018.01.0006

Midamegbe A., R. Vitalis, T. Malausa, E. Delava, S. Cros-Arteil, et al., 2011 Scanning the European corn borer (Ostrinia spp.) genome for adaptive divergence between host-affiliated sibling species. Mol. Ecol. 20: 1414–1430. https://doi.org/10.1111/j.1365-294X.2011.05035.x

Migeon A., E. Nouguier, and F. Dorkeld, 2010 Spider Mites Web: A comprehensive database for the Tetranychidae, pp. 557–560 in Trends in Acarology,.

Nouhaud P., J. Peccoud, F. Mahéo, L. Mieuzet, J. Jaquiéry, et al., 2014 Genomic regions repeatedly involved in divergence among plant-specialized pea aphid biotypes. J. Evol. Biol. 27: 2013–2020. https://doi.org/10.1111/jeb.12441

Oppenheim S. J., F. Gould, and K. R. Hopper, 2012 The genetic architecture of a complex ecological trait: host plant use in the specialist moth, Heliothis subflexa: the genetic architecture of host plant use. Evolution 66: 3336–3351. https://doi.org/10.1111/j.1558-5646.2012.01712.x

Oppenheim S. J., R. H. Baker, S. Simon, and R. DeSalle, 2015 We can’t all be supermodels: the value of comparative transcriptomics to the study of non-model insects: Comparative transcriptomics of non-model insects. Insect Molecular Biology 24: 139–154. https://doi.org/10.1111/imb.12154

R Core Team, 2016 R: A language and environment for statistical computing. R Foundation for Statistical Computing, Vienna, Austria.

Ranson H., M. G. Paton, B. Jensen, L. McCarroll, A. Vaughan, et al., 2004 Genetic mapping of genes conferring permethrin resistance in the malaria vector, Anopheles gambiae. Insect Mol. Biol. 13: 379–386. https://doi.org/10.1111/j.0962-1075.2004.00495.x

Riga M., A. Myridakis, D. Tsakireli, E. Morou, E. G. Stephanou, et al., 2015 Functional characterization of the Tetranychus urticae CYP392A11, a cytochrome P450 that hydroxylates the METI acaricides cyenopyrafen and fenpyroximate. Insect Biochemistry and Molecular Biology 65: 91–99. https://doi.org/10.1016/j.ibmb.2015.09.004

Riga M., S. Bajda, C. Themistokleous, S. Papadaki, M. Palzewicz, et al., 2017 The relative contribution of target-site mutations in complex acaricide resistant phenotypes as assessed by marker assisted backcrossing in Tetranychus urticae. Scientific Reports 7. https://doi.org/10.1038/s41598-017-09054-y

Robinson J. T., H. Thorvaldsdóttir, W. Winckler, M. Guttman, E. S. Lander, et al., 2011 Integrative genomics viewer. Nat. Biotechnol. 29: 24–26. https://doi.org/10.1038/nbt.1754

Roush R. T., and J. A. McKenzie, 1987 Ecological Genetics of Insecticide and Acaricide Resistance. Annual Review of Entomology 32: 361–380. https://doi.org/10.1146/annurev.en.32.010187.002045

Saavedra-Rodriguez K., C. Strode, A. F. Suarez, I. F. Salas, H. Ranson, et al., 2008 Quantitative Trait Loci Mapping of Genome Regions Controlling Permethrin Resistance in the Mosquito Aedes aegypti. Genetics 180: 1137–1152. https://doi.org/10.1534/genetics.108.087924

Schoonhoven L. M., J. J. A. van Loon, and M. Dicke, 2005 Insect-plant biology. Oxford University Press, Oxford; New York.

Shendure J., S. Balasubramanian, G. M. Church, W. Gilbert, J. Rogers, et al., 2017 DNA sequencing at 40: past, present and future. Nature 550: 345–353. https://doi.org/10.1038/nature24286

Snoeck S., N. Wybouw, T. Van Leeuwen, and W. Dermauw, 2018 Transcriptomic Plasticity in the Arthropod Generalist Tetranychus urticae Upon Long-Term Acclimation to Different Host Plants. G3&#58; Genes|Genomes|Genetics g3.200585.2018. https://doi.org/10.1534/g3.118.200585

Sparks T. C., and R. Nauen, 2015 IRAC: Mode of action classification and insecticide resistance management. Pestic Biochem Physiol 121: 122–128. https://doi.org/10.1016/j.pestbp.2014.11.014

Sterck L., K. Billiau, T. Abeel, P. Rouzé, and Y. Van de Peer, 2012 ORCAE: online resource for community annotation of eukaryotes. Nature Methods 9: 1041–1041. https://doi.org/10.1038/nmeth.2242

Strong D. R., J. H. Lawton, and T. R. E. Southwood, 1984 Insects on plants: community patterns and mechanisms. Blackwell, Oxford.

Thorvaldsdóttir H., J. T. Robinson, and J. P. Mesirov, 2013 Integrative Genomics Viewer (IGV): high-performance genomics data visualization and exploration. Bioinformatics 14: 178–192. https://doi.org/10.1093/bib/bbs017

Van der Auwera G. A., M. O. Carneiro, C. Hartl, R. Poplin, G. del Angel, et al., 2013 From FastQ data to high-confidence variant calls: The Genome Analysis Toolkit best practices pipeline, pp. 11.10.1-11.10.33 in Current Protocols in Bioinformatics, edited by Bateman A., Pearson W. R., Stein L. D., Stormo G. D., Yates J. R. John Wiley & Sons, Inc., Hoboken, NJ, USA.

Van Leeuwen T., B. Vanholme, S. Van Pottelberge, P. Van Nieuwenhuyse, R. Nauen, et al., 2008 Mitochondrial heteroplasmy and the evolution of insecticide resistance: Non-Mendelian inheritance in action. Proceedings of the National Academy of Sciences 105: 5980–5985. https://doi.org/10.1073/pnas.0802224105

Van Leeuwen T., J. Vontas, A. Tsagkarakou, W. Dermauw, and L. Tirry, 2010 Acaricide resistance mechanisms in the two-spotted spider mite Tetranychus urticae and other important Acari: A review. Insect Biochemistry and Molecular Biology 40: 563–572. https://doi.org/10.1016/j.ibmb.2010.05.008

Van Leeuwen T., P. Demaeght, E. J. Osborne, W. Dermauw, S. Gohlke, et al., 2012 Population bulk segregant mapping uncovers resistance mutations and the mode of action of a chitin synthesis inhibitor in arthropods. Proceedings of the National Academy of Sciences 109: 4407–4412. https://doi.org/10.1073/pnas.1200068109

Van Leeuwen T., and W. Dermauw, 2016 The molecular evolution of xenobiotic metabolism and resistance in chelicerate mites. Annual Review of Entomology 61: 475–498. https://doi.org/10.1146/annurev-ento-010715-023907

Van Pottelberge S., T. Van Leeuwen, J. Khajehali, and L. Tirry, 2009 Genetic and biochemical analysis of a laboratory-selected spirodiclofen-resistant strain of Tetranychus urticae Koch (Acari: Tetranychidae). Pest Management Science 65: 358–366. https://doi.org/10.1002/ps.1698

Weetman D., L. S. Djogbenou, and E. Lucas, 2018 Copy number variation (CNV) and insecticide resistance in mosquitoes: evolving knowledge or an evolving problem? Curr Opin Insect Sci 27: 82–88. https://doi.org/10.1016/j.cois.2018.04.005

Wickham H., 2016 ggplot2: Elegant Graphics for Data Analysis. Springer-Verlag New York, New York, NY.

Wybouw N., W. Dermauw, L. Tirry, C. Stevens, M. Grbić, et al., 2014 A gene horizontally transferred from bacteria protects arthropods from host plant cyanide poisoning. eLife 3:e02365. https://doi.org/10.7554/eLife.02365

Wybouw N., V. Zhurov, C. Martel, K. A. Bruinsma, F. Hendrickx, et al., 2015 Adaptation of a polyphagous herbivore to a novel host plant extensively shapes the transcriptome of herbivore and host. Mol. Ecol. 24: 4647–4663. https://doi.org/10.1111/mec.13330

Wybouw N., T. Van Leeuwen, and W. Dermauw, 2018 A massive incorporation of microbial genes into the genome of Tetranychus urticae, a polyphagous arthropod herbivore. Insect Mol. Biol. 27: 333–351. https://doi.org/10.1111/imb.12374

Zhao J.-Y., X.-T. Zhao, J.-T. Sun, L.-F. Zou, S.-X. Yang, et al., 2016 Transcriptome and proteome analyses reveal complex mechanisms of reproductive diapause in the two-spotted spider mite, Tetranychus urticae. Insect Molecular Biology. https://doi.org/10.1111/imb.12286

Zimmer C. T., W. T. Garrood, K. S. Singh, E. Randall, B. Lueke, et al., 2018 Neofunctionalization of Duplicated P450 Genes Drives the Evolution of Insecticide Resistance in the Brown Planthopper. Current Biology 28: 268-274.e5. https://doi.org/10.1016/j.cub.2017.11.060

